# Systematic revision and redefinition of the genus *Scirrotherium* Edmund and Theodor, 1997 (Cingulata, Pampatheriidae): Implications for the origin of pampatheriids and the evolution of the South American lineage including *Holmesina*

**DOI:** 10.1101/719153

**Authors:** Kevin Jiménez-Lara

## Abstract

The intrageneric relationships of the pampatheriid genus *Scirrotherium* and its affinities with supposedly related genera, i.e., *Kraglievichia* and *Holmesina*, are revised through parsimony phylogenetic analyses and new comparative morphological descriptions. For this work, unpublished material of pampatheriids (numerous osteoderms, one partial skull and a few postcranial bones) from Neogene formations of Colombia was analyzed. The results show that *Scirrotherium* is paraphyletic if we include all its referred species, i.e., *Scirrotherium hondaensis*, *S. carinatum* and *S. antelucanus*. The species *S. carinatum* is closer to *Kraglievichia paranensis* than to *S. hondaensis* or *S. antelucanus*, therefore the new name *K. carinatum* comb. nov. is proposed. The relationship among *S. hondaensis* and *S. antelucanus* could not be resolved, so these species should be designated in aphyly. In spite of failing to recover *S. hondaensis* and *S. antelucanus* as one single clade, here is preferred to maintain the generic name *Scirrotherium* in both species based on diagnostic evidence. New emended diagnoses for *Scirrotherium*, *S. hondaensis* and *Kraglievichia* are provided. The genus *Holmesina* was found to be monophyletic and positioned as the sister clade of *Scirrotherium* + *Kraglievichia*. The evolutionary and biogeographic implications of the new phylogeny and taxonomic re-arrangements are discussed. A possible geographic origin of the family Pampatheriidae and *Scirrotherium* in low latitudes of South America as early as Early Miocene times is claimed. The South American ancestor or sister taxon of *Holmesina* is predicted to be morphologically more similar to *Scirrotherium* than to *Kraglievichia*.

## 1. Introduction

The pampatheriids (Pampatheriidae) are a morphologically conservative extinct clade of glyptodontoid cingulates (Xenarthra: Glyptodontoidea *sensu* McKenna and Bell 1997) with medium-to-large body sizes (Edmund 1985; Góis et al. 2013). They were distributed from the Neogene to the Early Holocene in numerous localities in South America (their native range), Central America, Mexico and the United States (Edmund 1985; Vizcaíno et al. 1998; Rincón et al. 2014; Góis et al. 2015 and references therein). As the modern armadillos (Dasypodidae), pampatheriids have a flexible carapace characterized by the presence of three transverse bands of imbricated osteoderms, which form a kind of “articulation” between the scapular and pelvic shields (Edmund 1985). The pampatheriids also have multiple features, especially in their skull and mandible, which, collectively, define them as the sister group of glyptodontids –Glyptodontidae (Gaudin 2004; Gaudin and Wible 2006; Billet et al. 2011; Delsuc et al. 2012), namely, a deep horizontal mandibular ramus, a laterally-directed zygomatic root, a transversely wide glenoid fossa, rugose pterygoids, among others (Gaudin and Wible, 2006).

The fossil record of Pampatheriidae is mainly represented by isolated specimens, most of which are osteoderms and, to a lesser extent, skulls, mandibles and postcranial bones. Relatively complete and articulated skeletons are uncommon (Edmund 1985; Góis 2013). Due to this fact, the systematics of this group has historically been based on osteoderm characters (Edmund 1985, 1987; Góis et al. 2013), as has often been the case with other cingulate clades. Overall, nearly 20 pampatheriid species and seven genera are known (Góis 2013). The latter conform two possible subfamilial lineages: (1) that including to the genera *Plaina* and *Pampatherium*; and (2) that comprising the genera *Scirrotherium*, *Kraglievichia* and *Holmesina* (Edmund 1985). However, there is no any published phylogenetic analysis on the relationships among the different pampatheriid genera in the scientific literature. Only Góis (2013) performed a phylogenetic analysis of these taxa, but his results have not been published so far. In Góis’s consensus tree, it was corroborated the hypothesis on the two subfamilial lineages as suggested by Edmund (1985).

The genus *Scirrotherium* is the oldest undoubted pampatheriid (Góis et al. 2013; Rincón et al. 2014) and one of the four Miocene genera (the others are *Kraglievichia* Castellanos 1927; *Vassallia* Castellanos, 1927; and *Plaina* Castellanos, 1937). This taxon was originally described by Edmund and Theodor (1997) based on craniomandibular, postcranial and osteoderm specimens collected in the Middle Miocene (Serravalian) sedimentary sequence of the La Venta area, in southwestern Colombia. These authors suggested that the type and only known species in that time, *Scirrotherium hondaensis*, has plesiomorphic traits in its osteological morphology which are expected for its antiquity. Additionally, they highlighted the morphological similarity of *S. hondaensis* with the species *Vassallia minuta* (Late Miocene of southern and central South America; De Iullis and Edmund 2002), more than with any other pampatheriid.

Later, Góis et al. (2013) described a second species for *Scirrotherium*, *S. carinatum*, from the Late Miocene (Tortonian) of northeastern and southern Argentina and northwestern Brazil. In northeastern Argentina (Province of Entre Ríos), *S. carinatum* is found in the same basal stratigraphic levels (“Conglomerado Osífero”, literally meaning ‘bone-bearing conglomerate’) of the Ituzaingó Formation as the middle-sized pampatheriid *Kraglievichia paranensis* (Góis et al. 2013; Scillato-Yané et al. 2013), a taxon clearly distinct but not distantly related to *Scirrotherium*, as previously indicated. *Scirrotherium carinatum*, based exclusively on osteoderms from different regions of the armored carapace, has an estimated body size comparable or slightly smaller than that of *S. hondaensis* (Góis et al. 2013).

The phylogenetic analysis conducted by Góis (2013) recovered a polytomy involving *S. carinatum* and *S. hondaensis*, one of these species or both being the sister taxon/taxa of the clade *Kraglievichia* + *Holmesina* (except *H. floridanus*). Considering that *Scirrotherium* is the oldest known pampatheriid, it is notorious the non-basal position of the *Scirrotherium* species in the cladogram of Góis (2013). Instead, these species are closely allied with terminal taxa, i.e., *Holmesina* spp. (Edmund 1985, 1987; Gaudin and Lyon 2017). If this result is correct, it would indicate a significant ghost lineage at the base of Pampatheriidae.

Góis (2013) found phylogenetic support for *Scirrotherium* through one single synapomorphy, i.e., the presence of deep longitudinal depressions in the osteoderms. This is a distinctive feature of *S. carinatum* but the longitudinal depressions in *S. hondaensis* are relatively shallow. Interestingly, this putative synapomorphy is actually shared by *K. paranensis*. Góis (2013) explained the lack of phylogenetic resolution for *Scirrotherium* by noting the fragmentary character of the available fossil specimens of *S. hondaensis*. However, unlike the latter species, the skull, mandible and any postcranial bone of *S. carinatum* are unknown (Góis et al. 2013).

Laurito and Valerio (2013) reported new pampatheriid material from the Late Miocene (Tortonian to Messinian) of Costa Rica, which they assigned to a new species, *S. antelucanus*. This species, the largest referred to as *Scirrotherium* so far (body size comparable or slightly smaller than that of *K. paranensis*; Laurito and Valerio 2013), is based on osteoderms and some postcranial bones (femoral fragments and metatarsals). The occurrence of *S. antelucanus* in the Late Miocene of southern Central America suggests that *Scirrotherium* took part in the late Cenozoic biotic interchanges of the Americas earlier than any other pampatheriid (i.e., *Plaina*, *Pampatherium*, *Holmesina*; Woodburne 2010), invading tropical North America (“North America” is defined here as all the continental territories north of the ancient location of the main geographic barrier between the Americas during the early Neogene, i.e., the Central American Seaway in northwestern Colombia) before the definitive closing of the Panamanian Land Bridge (PLB) ca. 3 mya (Schmidt 2007; Coates and Stallard 2013; O’dea et al. 2016; Jaramillo 2018).

Recently, the occurrence of isolated osteoderms designated as *Scirrotherium* sp. or cf. *Scirrotherium* has been reported in several contributions on fossil vertebrate assemblages from the Neogene of Venezuela and Peru. On the basis of these discoveries, the geographic and chronological distribution of the genus has been expanded in such a way that this taxon is now known for the Early and Late Miocene (Burdigalian and Tortonian) of northwestern Venezuela (Rincón et al. 2014; Carrillo-Briceño et al. 2018) and Late Miocene (Tortonian) of eastern Peru (Antoine et al. 2016).

Assuming all the previous taxonomic assignments are correct, the latitudinal range of *Scirrotherium*, from southern Central America to Patagonia (southern Argentina), is the widest latitudinal range of a Miocene pampatheriid, comparable only with those of the Plio-Pleistocene forms *Pampatherium* and *Holmesina* (Scillato-Yané et al. 2005). This biogeographic evidence provides support for the hypothesis that *Scirrotherium* inhabited varied environments within its latitudinal range, and, consequently, that it probably had a relatively high ecological flexibility (Góis et al. 2013).

Despite the progress in the systematic and biogeographic research on *Scirrotherium*, a new reevaluation of several fundamental hypotheses about this taxon is needed, including its taxonomic definition and evolutionary relationships with other pampatheriid genera. Using parsimony phylogenetic analyses and comparative morphological descriptions of new pampatheriid remains from the Neogene of Colombia, this contribution reevaluates the taxonomic status of *Scirrotherium* and its relationships with supposedly allied genera, i.e., *Kraglievichia* and *Holmesina*. Accordingly, I suggest a new taxonomic and nomenclatural reorganization, with emended diagnoses for *Scirrotherium* and *Kraglievichia*.

Finally, considering the systematic reanalysis, I develop a model of biogeographic evolution for the lineage *Scirrotherium*-*Kraglievichia*-*Holmesina*. From this model, I draw out new hypotheses on the geographic origin of Pampatheriidae and the late Cenozoic dispersal events of pampatheriids to/from North America, including a possible re-entry event to South America for the species *S. antelucanus*.

## 2. Material and methods

### 2.1. Taxonomic sampling

I studied 12 species of pampatheriids attributed to six different genera. These species, in alphabetic order, are: *Holmesina floridanus* Robertson, 1976; *H. major* Lund, 1842; *H. occidentalis* Hoffstetter, 1952; *H. paulacoutoi* Cartelle and Bohórquez, 1985; *H. septentrionalis* Leidy, 1889; *Kraglievichia paranensis* Ameghino, 1888; *Pampatherium humboldtii* Lund, 1839; *Plaina intermedia* Ameghino, 1888; *Scirrotherium antelucanus* Laurito and Valerio, 2013; *S. carinatum* Góis, Scillato-Yané, Carlini and Guilherme, 2013; *S. hondaensis* Edmund and Theodor, 1997; and *Vassallia minuta* Moreno and Mercerat, 1891. Unidentified pampatheriid material (MUN STRI 16718 and 38064; see the section *Institutional abbreviations*) from the Castilletes Formation in Colombia (see below), which is referred to as “Castilletes specimens”, was also included in this selection.

Among the former nominal species, I follow Góis (2013) in considering *Vassallia maxima* as a junior synonym of *Pl. intermedia*. The only species of *Holmesina* not included in this study were *H. rondoniensis* Góis, Scillato-Yané, Carlini and Ubilla, 2012 and *H. cryptae* Moura, Góis, Galliari and Fernandes 2019. In the case of *H. rondoniensis*, its exclusion is based on a preliminary phylogenetic analysis in which it was identified as a “wildcard” taxon obscuring phylogenetic resolution as a result of lack of information on the osteoderm features of this species. On the other hand, the scientific article in which *H. cryptae* was described has been very recently published (Moura et al. 2019), after the completion of this work. Consequently, it was preferred to omit this species here. *Tonnicinctus mirus* Góis, González Ruiz, Scillato-Yané and Soibelzon, 2015 was also not included in this analysis given that this species is considered a highly divergent taxon without any apparent substantial interest with respect to the systematic issues here addressed.

### 2.2. Morphological description of the specimens

The osteological morphology of the selected species was re-examined from direct observations of specimens and published/unpublished descriptions (Simpson 1930, Castellanos 1937; Edmund 1985, 1987; Edmund and Theodor 1997; Góis 2013; Góis et al. 2013; Laurito and Valerio 2013; Scillato-Yané et al. 2013; Góis et al. 2015; Gaudin andg Lyon 2017). Naturally, according to the objectives of this research, during the revision of material I focused on the species *S. antelucanus*, *S. carinatum* and *S. hondaensis*, and, additionally, species of genera considered closely allied to *Scirrotherium*, i.e., *K. paranensis* and *Holmesina* spp. (particularly *H. floridanus*; Appendix S1 of the Supplementary Material).

On the other hand, new undescribed cranial, postcranial and osteoderm specimens were also used to reevaluate the morphological variability of *Scirrotherium*. This material comes from five Neogene geological units of Colombia (Fig. 1): (1) Castilletes Formation (Early to Middle Miocene, late Burdigalian-Langhian), Municipality of Uribia, Department of La Guajira; (2) La Victoria Formation (late Middle Miocene, Serravalian), Municipality of Villavieja, Department of Huila; (3) Villavieja Formation (late Middle Miocene, Serravalian), Municipality of Villavieja, Department of Huila; (4) Sincelejo Formation (Late Miocene-Early Pliocene, Messinian-Zanclean), Municipality of Los Palmitos, Department of Sucre; (5) Ware Formation (Late Pliocene, Piacenzian), Municipality of Uribia, Department of La Guajira. For detailed lithological descriptions and chronostratigraphic data on these formations, the reader is referred to the following references: Moreno et al. (2015) for the Castilletes and Ware Formations; Guerrero (1997), Flynn et al. (1997) and Anderson et al. (2016) for the La Victoria and Villavieja formations; and Flinch (2003), Villarroel and Clavijo (2005), Bermúdez et al. (2009) and Alfaro and Holz (2014) for the Sincelejo Formation. The new fossils are deposited in the Paleontological Collection of the Museo Mapuka de la Universidad del Norte, Barranquilla, Colombia, except those collected in the La Victoria and Villavieja Formations. The latter are housed at the Museo de Historia Natural La Tatacoa, La Victoria Town, Municipality of Villavieja, Department of Huila, Colombia.

**Figure 1.**
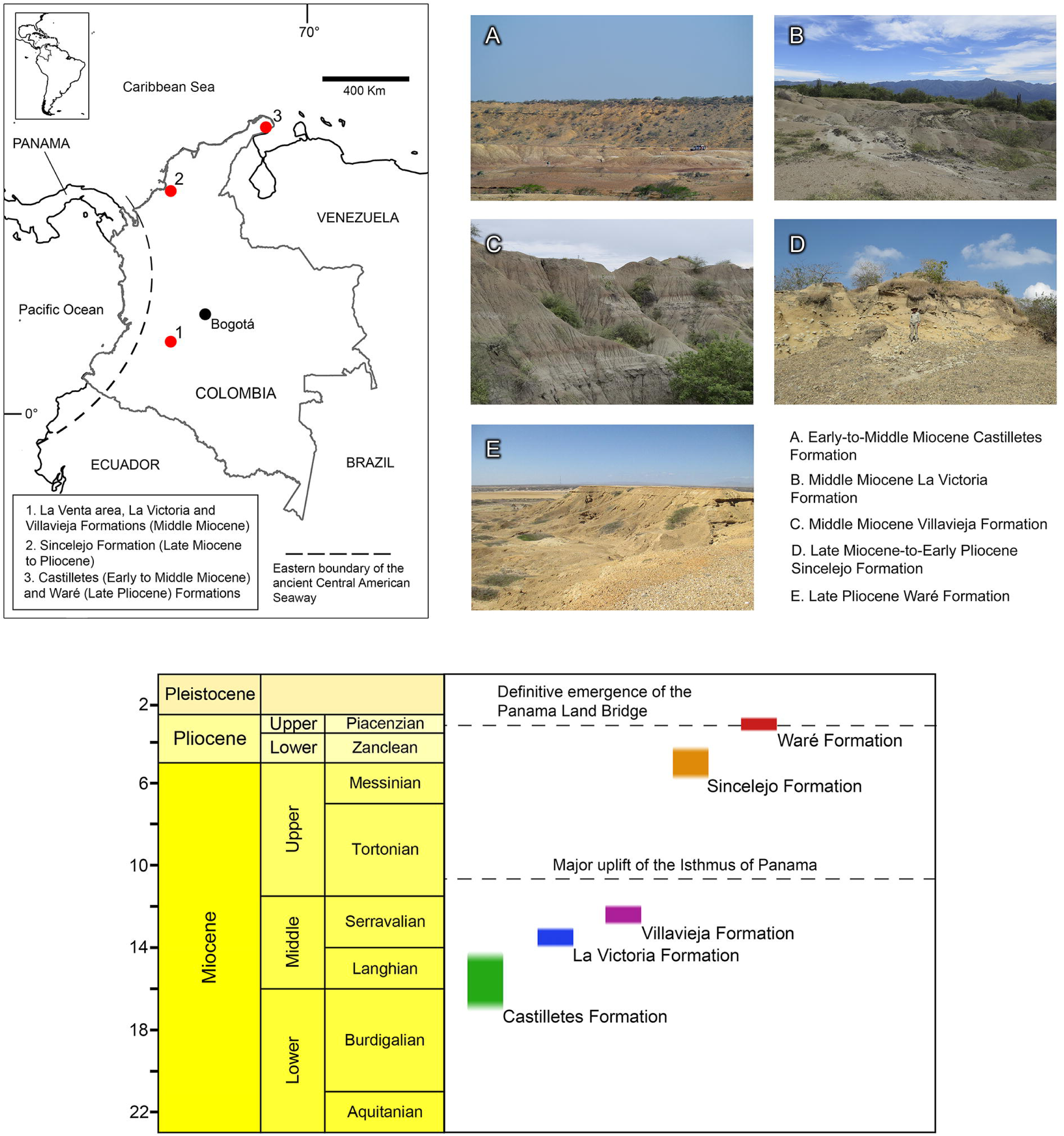
Geographic and stratigraphic provenance of the newly described material of pampatheriids from the Neogene of Colombia. In the left upper corner, a map of northwesternmost South America and the location of the regions of Colombia where there are outcrops of the formations with pampatheriid specimens for this study. In the right upper corner, photos of characteristic outcrops of these formations. Below in the center, a general chronostratigraphic scheme with the position of each formation within the Neogene and two important tectonic/palaeogeographic events in northwestern South America, i.e., a major, underwater uplift of the Isthmus of Panama (Schmidt 2007) and the definitive emergence of the Panamanian Land Bridge (O’dea et al. 2016). The photo of an outcrop of the Castilletes Formation was taken by Edwin Cadena.

Cranial measurements, all taken on the midline of the skull (dorsally or ventrally), follow Góis (2013). The anatomical terminology for osteoderms is based on the proposals by Góis et al. (2013). All the measurements were taken with a digital caliper.

### 2.3. Character matrix

I used exclusively cranial, dental and osteoderm characters, given that postcranial bones of most species of Pampatheriidae are poorly known (Góis 2013). The character construction was based on personal observations and previous quantitative and qualitative analyses of the interspecific, intergeneric and familial morphological variability of pampatheriids (e.g., Edmund 1985, 1987; Góis 2013; Góis et al. 2013; Laurito and Valerio 2013; Gaudin and Lyon, 2017). Overall, a matrix of 27 characters (Appendix S2 of the Supplementary Material) was built and managed on Mesquite version 2.75 (Maddison and Maddison 2010). If present, parsimony-uninformative characters were used to define potential autapomorphies of the taxa under study.

### 2.4. Cladistic analyses

Parsimony analyses under schemes of equal weights and implied weights (characters reweighted *a posteriori*; see below) were performed in PAUP* version 4.0a142 (Swofford 2015). In both weighting schemes, the species *P. humboldtii*, *Pl. intermedia* and *V. minuta* were selected as outgroup (sister group). This selection is based on the hypothesis about subfamilial relationships of Pampatheriidae by Edmund (1985) and the phylogeny of Góis (2013). All the characters were treated as unordered. The analyses were run with the branch and bound search to estimate maximum parsimony trees. The criterion for character optimization was DELTRAN (see Gaudin [2004] for justification of this configuration). For reordering of branches, the algorithm of tree-bisection-reconnection (TBR) branch swapping was used. The topological results of most parsimonious trees were summarized through strict consensus trees.

In this work, the methodology of implied weights was intended to mitigate potential biases arising from limited number of characters (especially osteoderm characters, as a consequence of the evolutionary trend in Pampatheriidae towards a simplification of the ornamentation in comparison with that in other cingulate clades, e.g., Glyptodontidae) and the effect of homoplastic characters (Goloboff et al. 2008; Goloboff 2014). Characters were reweighted using the rescaled consistency index (mean value) of the equally-weighted parsimony analysis (see Ausich et al. 2015 and references therein for justification of the use of rescaled consistency index for implied-weights parsimony analyses). A default concavity value (*k =* 3) was selected (Goloboff et al. 2018). Three successive rounds of character reweighting were needed until identical set of strict consensus trees were found in two consecutive searches (Swofford and Bell 2017).

Node stability for the strict consensus trees was evaluated using bootstrap resampling procedures (branch and bound search with 100 replicates). The software FigTree v1.4.3 (http://tree.bio.ed.ac.uk/software/figtree/) was used as a graphical viewer and editor for the cladograms.

### 2.5. Taxonomic and nomenclatural criteria

I applied a taxonomic and nomenclatural criterion reasonably, but not strictly, constrained by the phylogeny. This implies looking for a natural classification (i.e., based on monophyletic groups) without ignoring possible limitations of the phylogenetic inference related to the available information in the fossil record and major morphological gaps. Additionally, open nomenclature was used to indicate taxonomic uncertainty when necessary, following the general recommendations of Bengston (1988) and updated definitions by Sigovini et al. (2016) for the qualifiers of this semantic tool of taxonomy.

### 2.6. Institutional abbreviations

CFM, Museo Nacional de Costa Rica, Colección de fósiles de la sección de Geología, San José, Costa Rica; FMNH, Field Museum Natural History, Chicago, Illinois, USA; MACN, Museo Argentino de Ciencias Naturales “Bernardino Rivadavia”, Colección de Paleovertebrados, Ciudad Autónoma de Buenos Aires, Argentina; MCL, Museu de Ciências Naturais da Pontifícia Universidade Católica de Minas Gerais, Belo Horizonte, Brazil; MG-PV, Museo Provincial de Ciencias Naturales Dr. Ángel Gallardo, Rosario, Argentina; MHD-P, Museo Histórico Departamental de Artigas, Artigas, Uruguay; MLP, Museo de La Plata, La Plata, Argentina; MUN STRI, Museo Mapuka de la Universidad del Norte, Colección de paleontología, Barranquilla, Colombia; ROM, Royal Ontario Museum, Toronto, Canadá; UCMP, University of California Museum of Paleontology, Berkeley, California, USA; UF, Florida Museum of Natural History, Gainesville, Florida, USA; UZM, Universitets Zoologisk Museum, Copenhagen, Denmark; VPPLT, Museo de Historia Natural La Tatacoa, Colección de paleontología, La Victoria Town, Huila, Colombia.

### 2.7. Anatomical abbreviations

FL, frontal bone length; GFL, greatest femoral length; GSL, greatest skull length; LUR, length of the upper teeth row; Mf, upper molariform; mf, lower molariform; NL, nasal bone length; PAL, parietal bone length; PL, hard palate length; TTW, maximum width at the third trochanter of the femur; DW, maximum width of the femoral distal epiphysis.

## 3. Results

The parsimony analysis with equal weights obtained 107 most parsimonious trees (MPTs), each one of these with a tree length of 44 steps (consistency index = 0.909; retention index = 0.907; rescaled consistency index = 0.825). The strict consensus tree from these MPTs (Fig. 2 (A); tree length = 52; consistency index = 0.769; retention index = 0.721; rescaled consistency index = 0.555) is not fully resolved because it has two polytomies. One of these polytomies involves the species *S. hondaensis*, *S. antelucanus* and *H. floridanus*, whereas the other is formed by *H. septentrionalis*, *H. major*, *H. paulacoutoi* and *H. occidentalis*. Three clades were recovered (excluding that of the entire ingroup): (1) All the ingroup taxa except “Castilletes specimens”; (2) *S. carinatum* + *K. paranensis*; and (3) *Holmesina* spp. except *H. floridanus*. On the other hand, the parsimony analysis with implied weights yielded 30 most parsimonious trees (MPTs) with a tree length of 109 weighted steps (consistency index = 0.982; retention index = 0.982; rescaled consistency index = 0.964). The strict consensus tree from the MPTs (Fig. 2(B); tree length = 91; consistency index = 0.978; retention index = 0.980; rescaled consistency index = 0.959), like that produced by the equally weighted approach, is not fully resolved. Again, two polytomies resulted, but in this case the polytomy including *S. hondaensis*, *S. antelucanus* and *H. floridanus* was altered. The latter taxon was placed as the basal-most *Holmesina* species. The polytomy formed by *H. septentrionalis*, *H. major*, *H. paulacoutoi* and *H. occidentalis* was unmodified. As a consequence of the relocation of *H. floridanus* within the topology, four clades were recovered: (1) All the ingroup taxa except “Castilletes specimens”; (2) *S. carinatum* + *K. paranensis*; (3) *Holmesina* spp.; and (4) *H. septentrionalis*, *H. major* + *H. paulacoutoi* + *H. occidentalis*. According to the two schemes of weighting for the parsimony analyses, *Scirrotherium* is paraphyletic if it is comprised of *S. antelucanus*, *S. hondaensis* and *S. carinatum*. *Scirrotherium carinatum* is closer to *K. paranensis* than to *S. hondaensis* or *S. antelucanus*. The relationship among *S. hondaensis* and *S. antelucanus* is not resolved in either of the two strict consensus trees.

**Figure 2.**
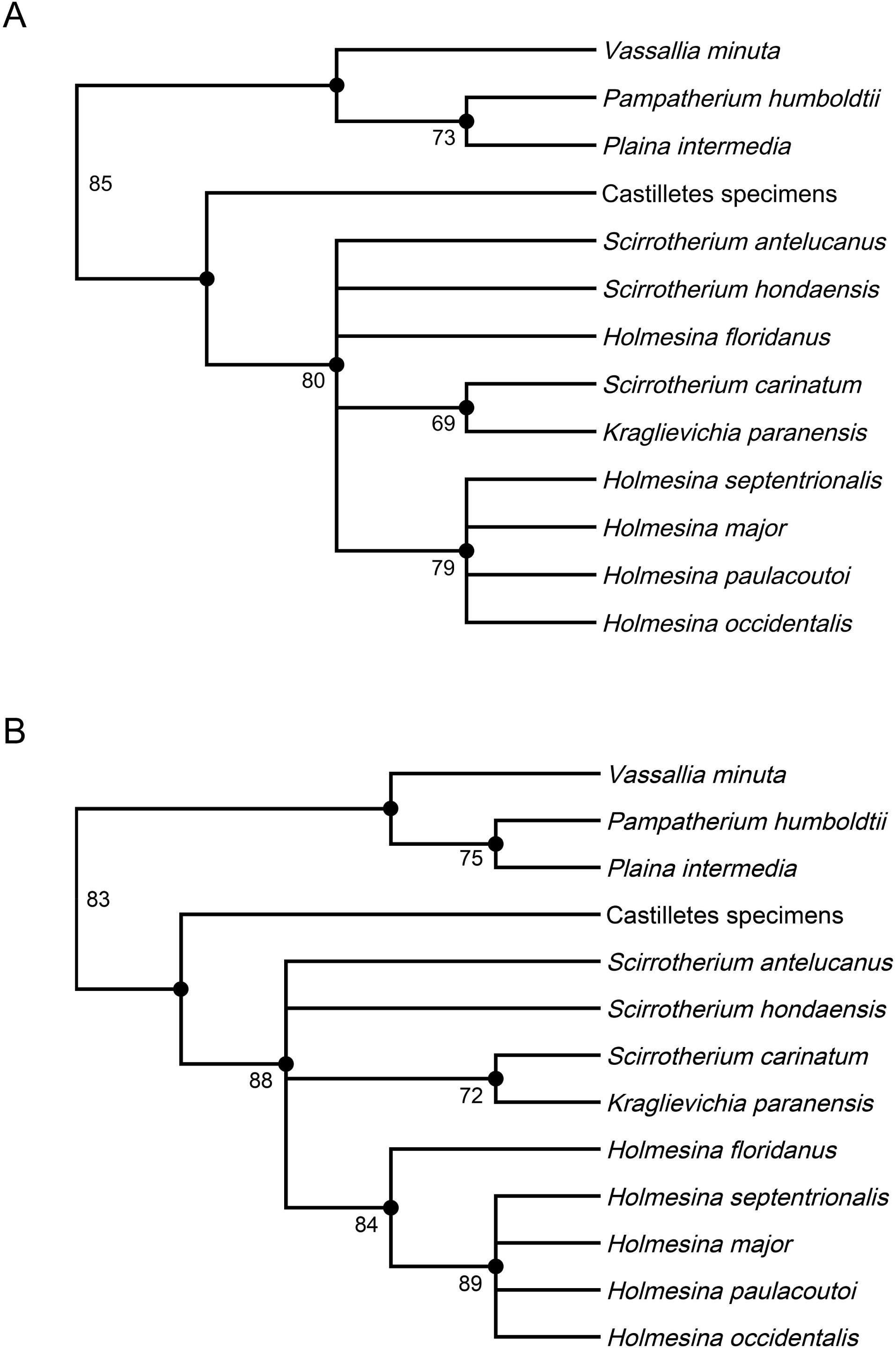
Phylogenetic results. A, strict consensus tree of the parsimony with equal weights. B, strict consensus tree of the parsimony analysis with implied weights. The numbers below nodes are bootstrap resampling frequencies. Note the difference in the phylogenetic position of *Holmesina floridanus* in the two strict consensus trees. Explanation of this difference in the *Discussion* section.

Under the equal weights analysis, all the nodes that include taxa of interest show (nearly) significant stability (resampling frequencies ca. 70 or greater). As expected, a similar result was obtained in the bootstrap resampling under implied weights, but with improved frequencies for nearly all the branches (except the basal-most branch, whose frequency decreased slightly from that of the former bootstrapping [85 to 83]).

## 4. **Systematic paleontology**

Xenarthra Cope, 1889

Cingulata Illiger, 1811

Glyptodontoidea Gray, 1869

Pampatheriidae Paula Couto, 1954

*Scirrotherium* Edmund and Theodor, 1997

*LSID.* urn:lsid:zoobank.org:act:313358B5-3B1F-4902-8C2E-BB07CFCBEE18

**Type species**: *Scirrotherium hondaensis* Edmund and Theodor, 1997 by original designation.

**Emended diagnosis**: A pampatheriid of small-to-middle body size that can be distinguished from other pampatheriids by the following combination of features: thin non-marginal fixed osteoderms (ca. 3.5–7 mm in thickness); slightly to moderately rough external surface of osteoderms; external surface of osteoderms with a sharp and uniformly narrow longitudinal central elevation; longitudinal central elevation from superficial to well-elevated; (very) shallow longitudinal depressions with gentle slope towards the marginal elevations; usually one single, transversely elongated row of large foramina in the anterior margin of fixed osteoderms; maximum number of foramina per row between 6 and 11.

**Remarks**: The taxonomic status of *Scirrotherium* is saved from invalidity by paraphyly by exclusion of the species ‘*S.*’ *carinatum* from the genus (see below). However, according to the preferred phylogenetic hypothesis presented here, i.e., the strict consensus tree from the parsimony analysis under implied weights (Fig. 2(B)), the other two referred species of *Scirrotherium*, *S. antelucanus* and *S. hondaensis*, should be designated in aphyly because they do not have resolved relationship between them (see Ebach and Williams 2010 for details about the phylogenetic concept of aphyly). Until new evidence becomes available, maintenance of the taxonomic validity of *Scirrotherium*, as defined here, is based on the emended diagnosis of this taxon, which is partially built from ambiguous synapomorphies, as well as from qualitative and quantitative morphological differences with other genera.

***Scirrotherium hondaensis*** Edmund and Theodor, 1997

*LSID.* urn:lsid:zoobank.org:act:E3B83181-91D6-44C8-90C0-BBAACEC2CDEE

**Holotype**: UCMP 40201, incomplete skull and left hemimandible (Edmund and Theodor, 1997).

**Type locality and horizon**: Municipality of Villavieja, Department of Huila, Colombia. La Victoria Formation, upper Middle Miocene, Serravalian (Edmund and Theodor, 1997).

**Emended differential diagnosis.** Pampatheriid of small body size that differs from other pampatheriids based on this unique combination of characters: external surface of osteoderms with ornamentation (especially the longitudinal central elevation and marginal elevations), in general terms, more protuberant than in *S. antelucanus*, but less than in *Kraglievichia*; size range of fixed osteoderms smaller than in *S. antelucanus* and similar to that in *Kraglievichia carinatum* comb. nov. (= ‘*S.*’ *carinatum*; Góis et al. 2013; see below); fixed osteoderms generally thicker than in *K. carinatum* comb. nov. but less than in *K. paranensis*, similar to *S. antelucanus*; anterior foramina smaller than in *S. antelucanus*; anterior foramina in fixed osteoderms usually aligned in one individual row, although infrequently these osteoderms show an extra, short or reduced row of anterior foramina; two rows of anterior foramina in mobile osteoderms, similar to *Vassallia* (Góis 2013); mf9 incipiently bilobed; frontals prominently convex in lateral view, with this convexity positioned posterior to the insertion of the anterior root of the zygomatic arch; anterior root of the zygomatic arch posterolaterally projected with respect to the main body of maxilla.

**Referred material**: VPPLT 004, several fixed osteoderms; VPPLT 264, several fixed osteoderms and one semi-mobile osteoderm; VPPLT 348, tens of fixed and (semi) mobile osteoderms; VPPLT 701, several fixed osteoderms; VPPLT 706, one anterior skull, one femoral diaphysis, one ulna without distal epiphysis, several vertebrae and numerous fixed and (semi) mobile osteoderms; VPPLT 1683 - MT 18, several fixed and (semi) mobile osteoderms; UCMP 39846, one proximal femoral epiphysis, one left calcaneum and one left astragalus. All the osteoderms referred to as *S. hondaensis* are illustrated in Fig. 3. Other important specimens are illustrated in Figs. 4–7 and 9(C).

**Figure 3.**
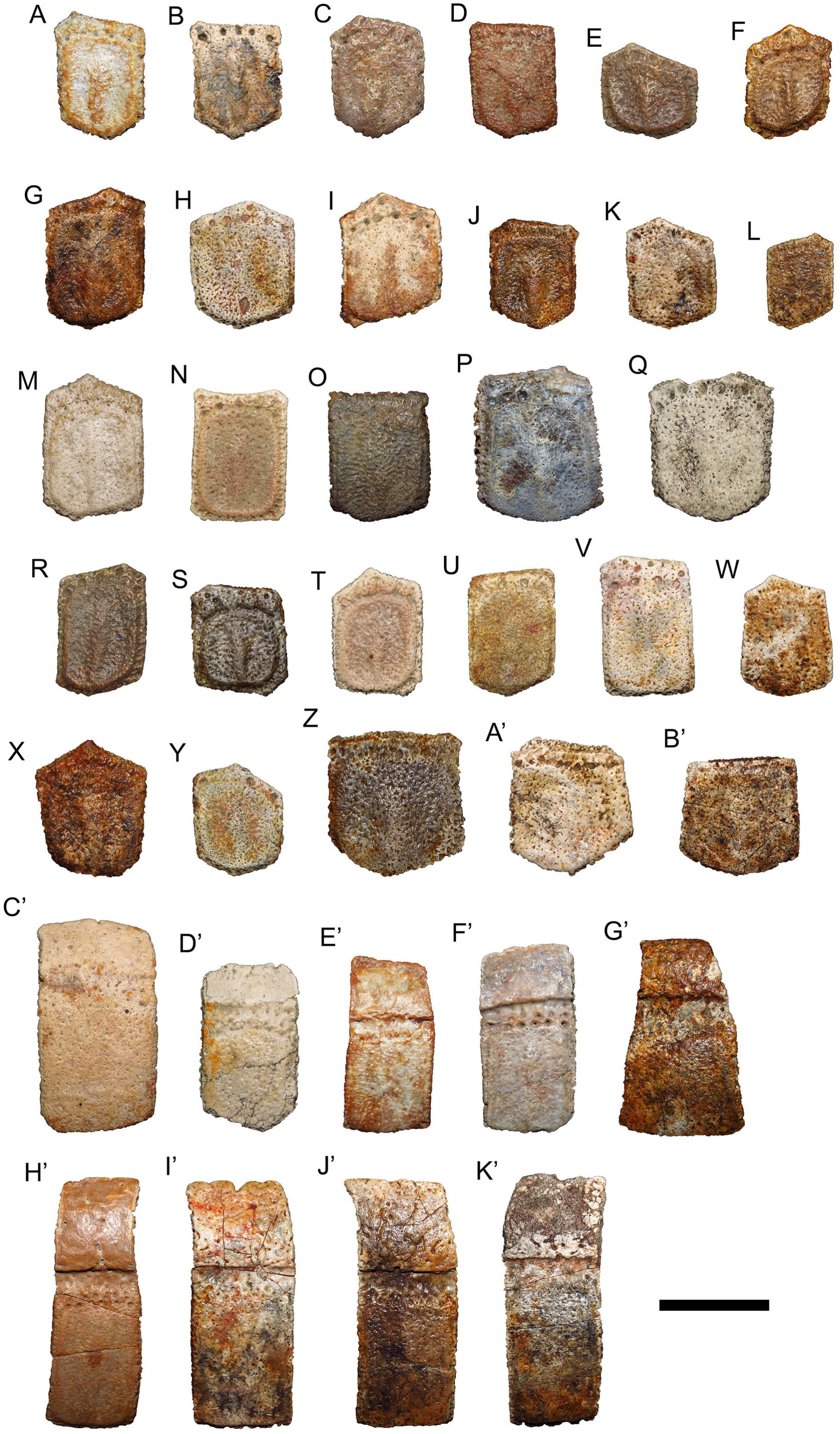
Fixed and (semi) mobile osteoderms of *Scirrotherium hondaensis* from the La Victoria and Villavieja Formations, Municipality of Villavieja, Department of Huila, Colombia. A–B’, fixed osteoderms; C’–K’, (semi) mobile osteoderms. The osteoderms G, J, K, L, W, X, Y, Z, A’, B’, G’, I’, J’ and K’ are associated with the catalog number VPPLT 348. The osteoderms H, U and V are associated with the catalog number VPPLT 004. The osteoderms T and D’ are associated with the catalog number VPPLT 701. All the former osteoderms come from the lower and middle La Victoria Formation. The osteoderms B, C, F, I, O, P, S, C’ and F’ are associated with the catalog number VPPLT 1683 - MT 18 and come from the top of the La Victoria Formation. The osteoderms A, D, E, M, N, Q, R, E’ and H’ are associated with the catalog number VPPLT 1683 - MT 18 and come from the lower Villavieja Formation. Scale bar equal to 20 mm.

**Occurrence**: VPPLT 004, 264, 701, 706 and (partially) 1683 - MT 18 were collected in the La Victoria Formation, upper Middle Miocene (Serravalian; see Figs. 3–6 for more details on the stratigraphic provenance of individual specimens), while the UCMP 39846 and part of VPPLT 1683 - MT18 comes from the Villavieja Formation, upper Middle Miocene (Serravalian).

**Description**: For the original and detailed description of this species, including its osteoderms, see Edmund and Theodor (1997). See the Tables 1 and 2 for an updated compilation of osteoderm measurements of the *Scirrotherium* species and comparisons with those of related taxa. Below there are descriptions of osteological structures and traits incompletely known or unknown for *S. hondaensis* so far.

**Table 1.** Fixed (scapular and pelvic) osteoderm measurements for taxa of interest in this study.

| Taxon/Measurement | Length | Width | Thickness | References |
| --- | --- | --- | --- | --- |
| *S. hondaensis* | 16-35.2 | 17.5-27.9 | 3.7-6.9 | This work; Góis et al. 2013 |
| *S. antelucanus* | 28.6-40.9 | 22-32.4 | 4.9-7.1 | This work; Laurito and Valerio 2013 |
| *K. carinatum* comb. nov. | 20.9-33.5 | 17-26.1 | 4.1-5.9 | This work; Góis et al. 2013 |
| *K. paranensis* | 30-45 | 22.5-28.3 | 6-11 | Góis et al. 2013 |
| *H. floridanus* | 24.4-36.7 | 18.9-32.1 | 6-9.7 | This work; Edmund 1987 |

**Table 2.** Mobile and semi-mobile osteoderm measurements for taxa of interest in this study.

| Taxon/Measurement | Length | Width | Thickness | References |
| --- | --- | --- | --- | --- |
| *S. hondaensis* | 29.4-60 | 17.9-27.4 | 4.9-7.3 | This work; Góis et al. 2013 |
| *S. antelucanus* | 38.2-64.6 | 19.4-28.9 | - | Laurito and Valerio 2013 |
| *K. carinatum* comb. nov. | 32-54.5 | 17-28.9 | 3.9-6 | This work; Góis et al. 2013 |
| *K. paranensis* | 60.5-70.5 | 25-29 | 7-9 | Góis et al. 2013 |
| *H. floridanus* | 61.8-71 | 17.6-28.5 | 4.7-6.3 | This work; Edmund 1987 |

*Skull*: The holotype of *S. hondaensis* UCMP 40201 includes a very fragmentary skull. This specimen does not preserve the anterior end of the rostrum, much of the orbit (both dorsally and ventrally), part of the upper dental series, ear region, braincase and occipital region. In comparison, the skull VPPLT 706 (Fig. 4) described here, is more complete, despite the fact that it is also missing some structures. This new, small skull (see Table 3 for morphometric comparisons) is relatively well preserved from the orbit to the anterior end of rostrum, except for the anterior zygomatic arch and nasal bones. It also has a less deformed rostrum than the holotypic skull. The general aspect of the new skull is similar to those of other pampatheriids. In lateral view, it is markedly depressed towards its anterior end. In dorsal view, it is also tapered towards its anterior tip, where it ends abruptly (Castellanos, 1937). Proportionally, the rostrum is shorter than that of *K. paranensis* and even more than that of *H. floridanus*. In lateral view, the facial process of premaxilla is less well defined than that of *H. floridanus*, although the premaxilla-maxilla suture has a convex form like the latter species (Gaudin and Lyon, 2017). The antorbital fossa is arranged more vertically than those of *K. paranensis* and *H. floridanus*. The lacrimal is, proportionally, the largest among pampatheriids. This bone precludes the frontomaxillary contact (restricted contact in the skull of *K. paranensis*). The dorsal contribution of the lacrimal to the orbit is, proportionally, greater than in *H. floridanus* but similar to that in *K. paranensis*. The anterior root of zygomatic arch is projected posterolaterally, unlike other pampatheriids whose skull is known, where the anterior root projects laterally. The frontals show a conspicuous convexity in a position posterior to the insertion of the anterior root of zygomatic arch. In dorsal view, the frontals are more anteroposteriorly elongated and more laterally expanded than in *K. paranensis*, but are similar to those of *Holmesina* spp. In ventral view, the hard palate has a wide aspect, since the rostrum is shortened in comparison with other pampatheriids. Only two anterior molariforms are preserved (the left Mf1 and the right Mf2), so inferences about upper dentition are made from the alveoli. The upper dental series, as in all the members of the family, is comprised of by nine molariforms. Of these, the last five (Mf5-Mf9) are bilobed. The anterior molariforms (Mf1-Mf4) converge anteriorly, but do not imbricate. The latter teeth are rounded to elliptical, similarly to the condition observed in *H. floridanus*. They also are less mesiodistally elongated than in *K. paranensis*. The molariforms with greatest occlusal area are the Mf5 and Mf6. The occlusal area of the upper molariforms decrease distally from the fifth and sixth molariforms to the ninth, as in all the pampatheriids. The Mf9 is the smallest of lobed molariforms and has the least degree of lobulation (elliptical shape for this tooth in the type material of *S. hondaensis*, according to Edmund and Theodor 1997). In ventral view, VPPLT 706 is characterized by a gradual transverse widening of the palatal portion of the maxilla from the level of the anterior border of the Mf5. The anterior portion of the palatines is preserved up to a level slightly posterior to the Mf9. The maxilla-palatine suture is not recognizable.

**Figure 4.**
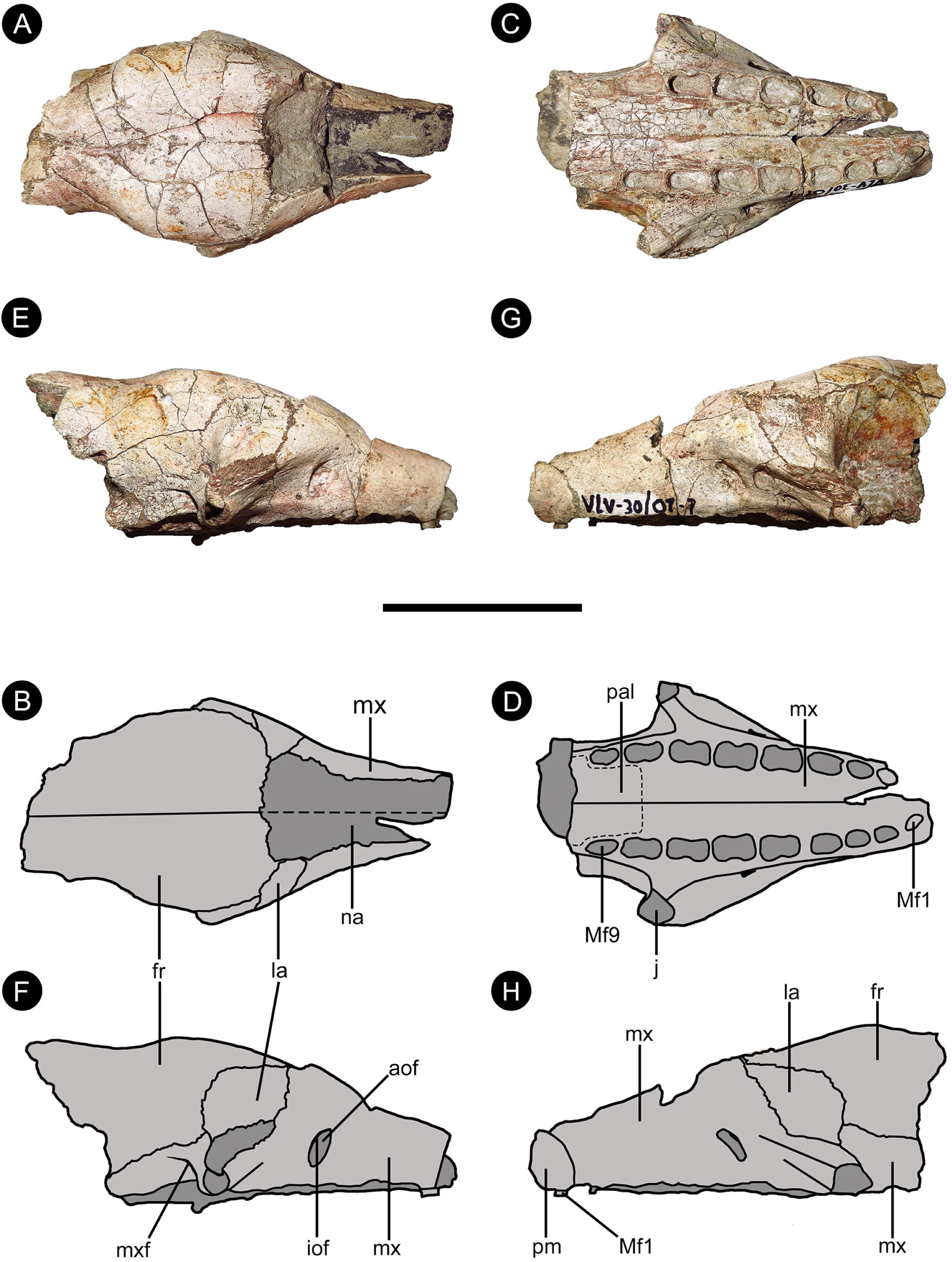
Photos and anatomical line drawings of the skull VPPLT 706 of *Scirrotherium hondaensis* from the middle La Victoria Formation, Municipality of Villavieja, Department of Huila, Colombia. A–B, dorsal views; C–D, ventral views; E–F, right lateral views; D–H, left lateral views. *Abbreviations*: aof, antorbital fossa; fr, frontal; iof, infraorbital foramen; la, lacrimal; Mf1, first upper molariform; Mf9, ninth upper molariform; mx, maxilla; mxf, maxillary foramen; na, nasal; pal, palatine; pm, premaxilla. Scale bar equal to 50 mm.

**Table 3.** Selected cranial measurements for VPPLT 706 of *Scirrotherium hondaensis* and related taxa whose skulls are known.

| Taxon/Measurement | GSL | NL | FL | PAL | LUR | PL | References |
| --- | --- | --- | --- | --- | --- | --- | --- |
| *S. hondaensis* | 117.3* | ~52.8 | ~55 | - | 84.1 | 94.3 | This work |
| *K. cf. paranensis* | 194 | 58 | 62 | 74 | - | 159 | This work |
| *H. floridanus* ** | 249 | 106.3 | 75 | 58.6 | 133.6 | 185 | This work |
| *H. septentrionalis* | 290 | - | - | - | 165 | 220 | Góis et al.<br>2012 |

\*Incomplete
\*\*Specimen UF 191448

*Femur*: This bone in *S. hondaensis* was largely unknown so far, except for a pair of epiphyses (proximal and distal) from a left femur in the UCMP collections (UCMP 39846). VPPLT 706 preserves a left femur (Fig. 5(A–D)) without epiphyses (apparently it is not the same bone from which the previously noted epiphyses came). Thus, the description of all these anatomical elements allows for a reconstruction of most aspects of femoral anatomy. The estimated proximo-distal length of this bone is ca. 162 mm, and its transverse width at the third trochanter is 27.6 mm. These morphometric values are the smallest known for femora of Pampatheriidae (Table 4). They are comparable only to those of MLP 69-IX-8-13A which was referred to as *K.* cf. *paranensis* (Góis 2013; Scillato-Yané et al. 2013). The femoral head is hemispheric and the greater trochanter is less high than that of *K.* cf. *paranensis*, but similar to the condition observed in *H. floridanus*. However, the greater trochanter has a more tapered proximal end than in the latter species. In *S. hondaensis*, the lesser trochanter is less mediolaterally expanded than in *K.* cf. *paranensis*. The femoral diaphysis is less curved mediolaterally than in *K.* cf. *paranensis*, similar to that of *H. floridanus*. The laterodistal border of the femur is more curved than in *H. floridanus*, similar to that of *K.* cf. *paranensis*. The third trochanter is, proportionally, larger than that of *K.* cf. *paranensis*, but is smaller than that of *H. floridanus*. The patellar facets are less defined or delimited than those of *K.* cf. *paranensis*. In *S. hondaensis* these facets are oriented toward the center of the anterior surface of the distal epiphysis, rather than laterally as in *K.* cf. *paranensis* and *H. floridanus*.

**Figure 5.**
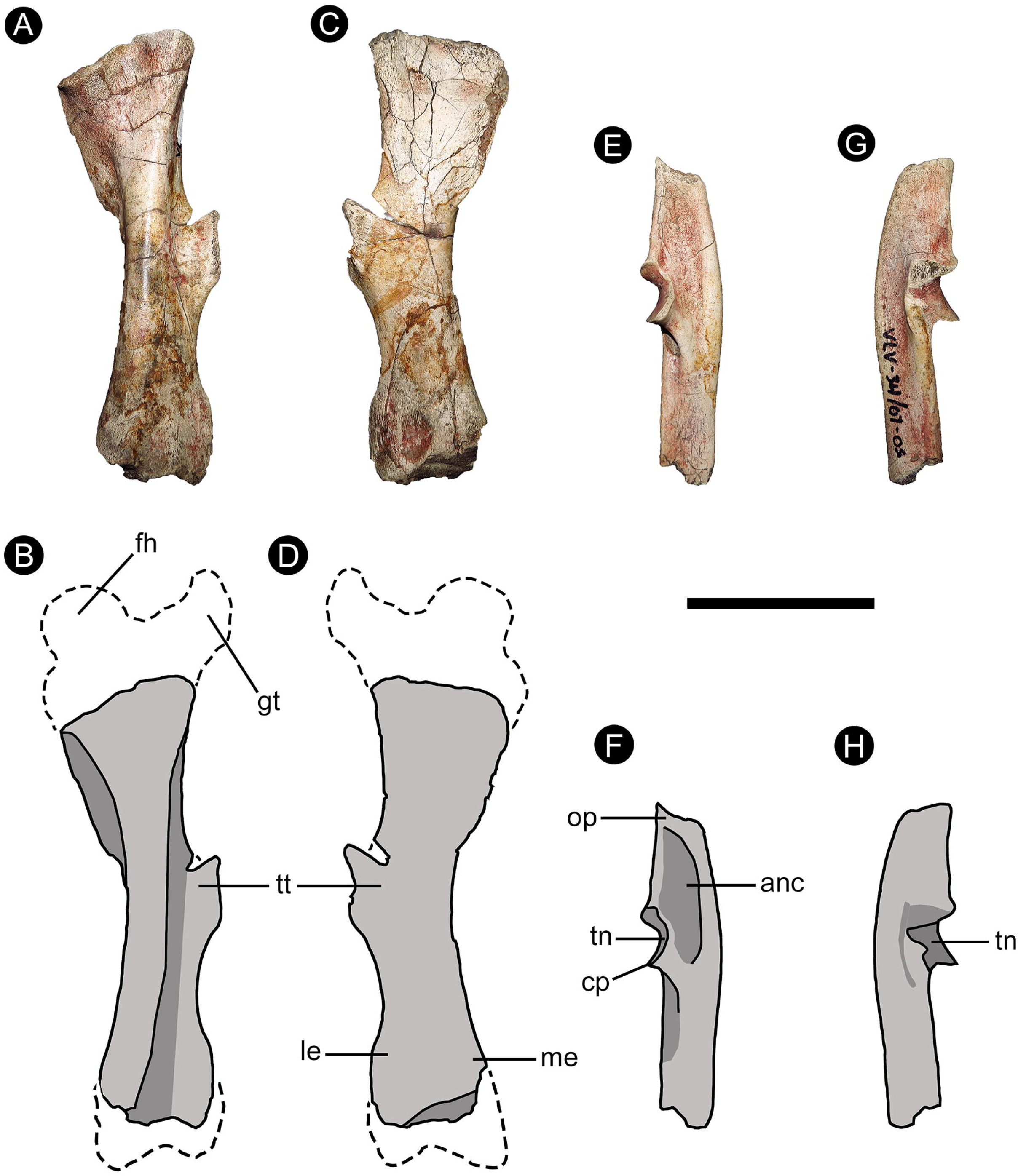
Photos and anatomical line drawings of the left femur and right ulna VPPLT 706 of *Scirrotherium hondaensis* from the middle La Victoria Formation, Municipality of Villavieja, Department of Huila, Colombia. The epiphyses of this femoral diaphysis have been reconstructed from those with catalog number UCMP 39846. A–B, anterior views of the femur; C–D, posterior views of the femur. E–F, medial views of the ulna; G–H, lateral views of the ulna. *Abbreviations*: anc, fossa for the anconeus muscle; cp, coronoid process; fh, femoral head; gt, greater trochanter; le, lateral epicondyle; me, medial epicondyle; op, olecranon process; tn, trochlear notch; tt, third trochanter. Scale bar equal to 50 mm.

**Table 4.** Femoral measurements for *Scirrotherium hondaensis* and related taxa whose femur is known.

| Measurement/Taxon | GFL | TTW | DW | References |
| --- | --- | --- | --- | --- |
| *S. hondaensis* | 162* | 27.6 | 32.5 | This work |
| *K. cf. paranensis* | 164 | 33.7 | 38 | This work |
| *H. floridanus* ** | 195 | 41 | 47 | This work |
| *H. septentrionalis* | 290 | 70 | 86 | Góis 2013 |

\*Estimated from the specimens VPPLT 706 and UCMP 39846
\*\*Specimen UF 24918

*Ulna*: This bone is described here for the first time in *S. hondaensis*. A right ulna (Fig. 5(E–H)) missing part of the diaphysis and the distal epyphysis is preserved in VPPLT 706. The estimated proximo-distal length is 89.5 mm. The olecranon is elongated and protuberant. In medial view, it is less proximally tapered than that of *H. floridanus*. The lateral entrance to the trochlear notch is very restricted, similarly to *Holmesina*. Likewise, it is less proximo-distally elongated than that of *H. floridanus*. Proximally, at the level of the trochlear notch, the posterior border is uniformly convex, not slightly concave as it is in *H. floridanus*. The depression for the insertion of the anconeus muscle is deep and proximally located, like that of *Holmesina*.

*Vertebrae*: Several vertebrae are also preserved in VPPLT 706 (Fig. 6). One of these is a thoracic vertebra and five are caudal vertebrae, of which four are articulated in two pairs and one is an isolated distal caudal vertebra. The body of the thoracic vertebra is eroded anteriorly, as are the anterior zygapophyses. Posteriorly, the vertebral body has an outline similar to that of other pampatheriids. Notably, two ventrolateral apophyses are projected from the vertebral body. Although fragmented, the neural spine of the same vertebra appears to be proportionally shorter than in *H. floridanus*. The anterior caudal vertebrae have a posteriorly oriented and tall neural spine. The transverse processes are relatively short.

**Figure 6.**
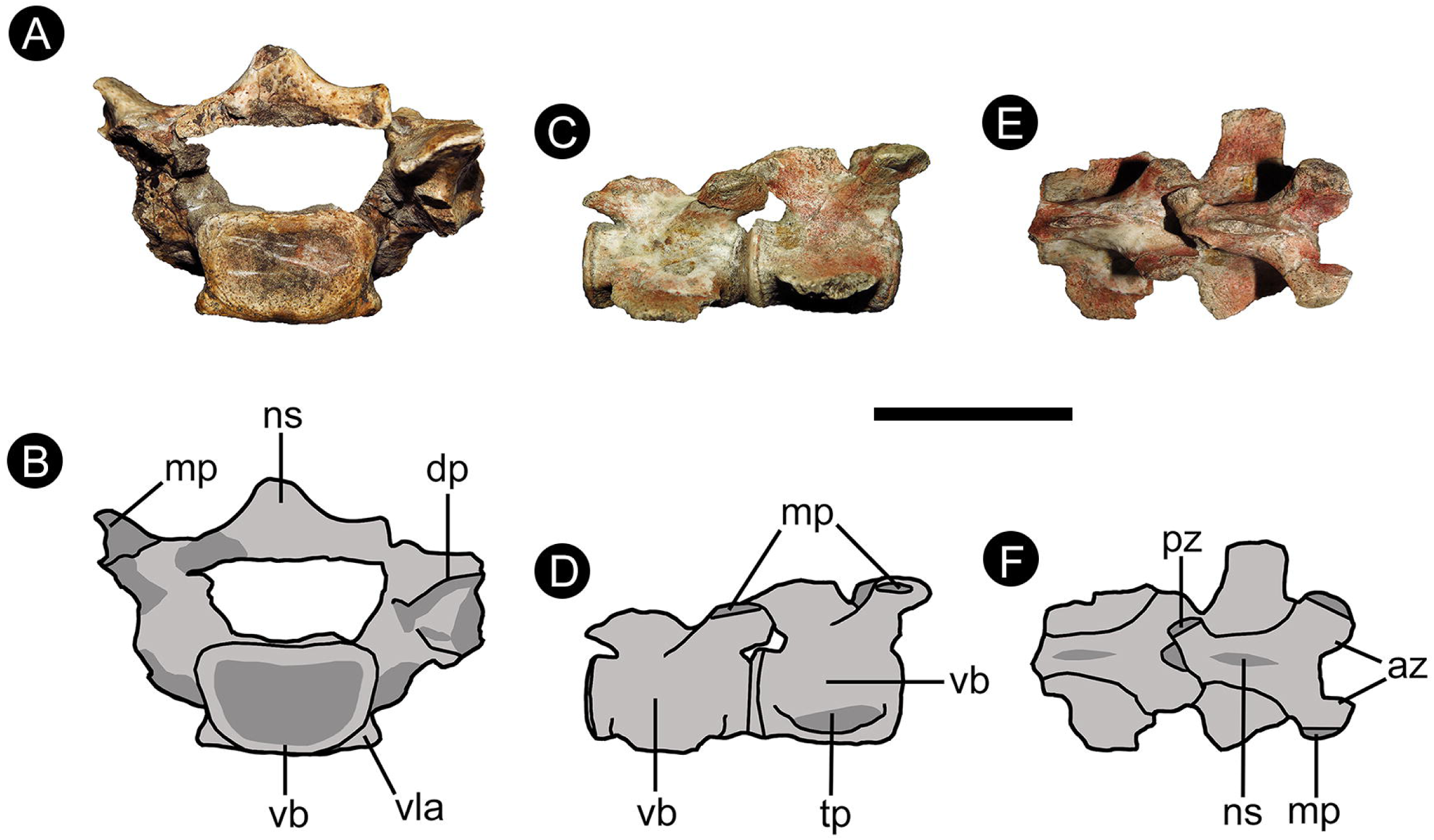
Photos and anatomical line drawings of a thoracic vertebra (A–B) and several anterior caudal vertebrae (C–F) VPPLT 706 of *Scirrotherium hondaensis* from the middle La Victoria Formation, Municipality of Villavieja, Department of Huila, Colombia. A–B, posterior views of the thoracic vertebra. C–D, lateral views of caudal vertebrae; E–F, dorsal views of caudal vertebrae. *Abbreviations*: az, anterior zygapophyses; mp, metapophyses; ns, neural spine; tp, transverse processes; vb, vertebral body; vla, ventrolateral apophyses. Scale bar equal to 30 mm.

*Astragalus*: Edmund and Theodor (1997) mentioned the existence of numerous undetermined postcranial elements whose description was to be postponed to a subsequent publication. However, that description was never published. This postcranial material from the UCMP collections includes a left astragalus (UCMP 39846; Fig. 7(A–D)). In dorsal view, the lateral trochlea is considerably larger than the medial trochlea. The astragalar head is bulging, spherical, and almost uniformly convex. There is a shallow concavity in the dorsal margin of the astragalar head whose function has been not determined, but it could be for tendinous insertion. The astragalar neck is well-differentiated, similar to that in *Holmesina*. In ventral view, the facets of articulation with the calcaneum, i.e., ectal and sustentacular, are widely separated, as one would expect by observing their counterparts on the calcaneum (Edmund 1987). The ectal facet is noticeably larger than the sustentacular facet, in contrast to the condition in *H. floridanus*. The ectal facet is kidney-shaped and the sustentacular facet has a sub-oval shape. Both of them are concave, especially the ectal facet, which is very deep. The sustentacular facet is located in a central position within the astragalar neck.

**Figure 7.**
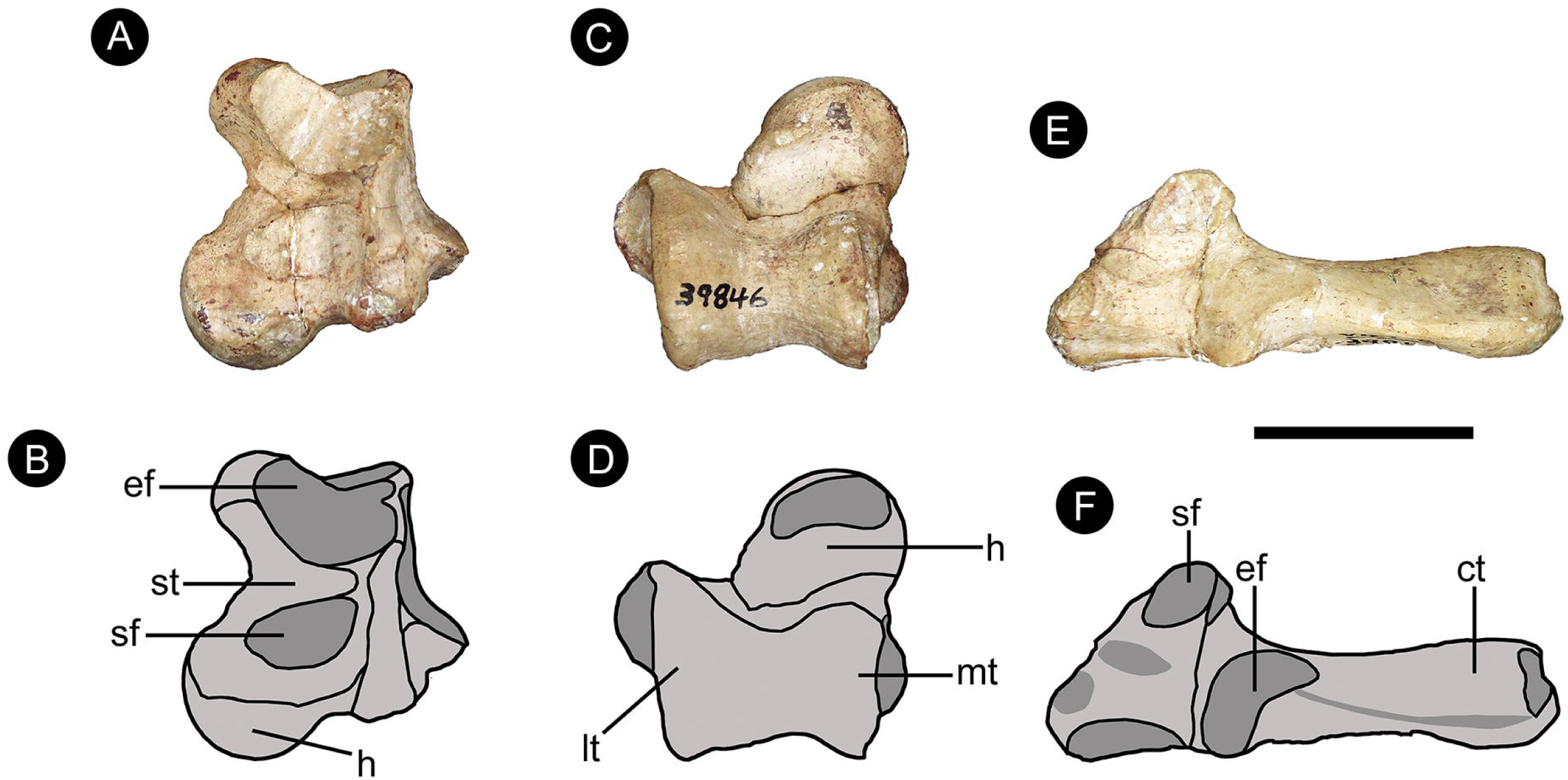
Photos and anatomical line drawings of the astragalus (A–D) and calcaneum (E – F) UCMP 39846 of *Scirrotherium hondaensis* from the lower (?) Villavieja Formation, Municipality of Villavieja, Department of Huila, Colombia. A–B, astragalus in plantar views; C–D, astragalus in dorsal views. E–F, calcaneum in dorsal views. *Abbreviations*: ct, calcaneal tuber; ef, ectal facet; h, head of the astragalus; lt, lateral trochlea; mt, medial trochlea; sf, sustentacular facet; st, sulcus tali. Scale bar equal to 20 mm.

*Calcaneum*: This bone is other postcranial element not described by Edmund and Theodor (1997) for *S. hondaensis*. UCMP 39846 is a well-preserved left calcaneum (Fig. 7(E–F)). It has proximo-distal length of 54.12 mm and a width at the level of facets (ectal and sustentacular) of ca. 10.2 mm. These values are the smallest known for calcanei referred to as Pampatheriidae. The only species whose calcaneum is comparable in size to that of *S. hondaensis* is *H. floridanus*. The calcaneum of the latter species is slightly more proximo-distally elongated than in *S. hondaensis*, but it is roughly two times wider at the level of the ectal and sustentacular facets. This means that the calcaneum of *H. floridanus* is more robust than that of *S. hondaensis*. The calcaneal head is anteroposteriorly elongated, like in *H. floridanus* and unlike the proportionally short calcaneal head of *H. septentrionalis*. The anterior end of the calcaneal head is less shortened than that of *Holmesina*. The calcaneum of *S. hondaensis* shows no contact between the ectal and sustentacular facets, similar to the condition in species of *Holmesina* other than *H. floridanus* (Góis 2013). These facets are slightly convex and they are separated by a moderately deep and very wide groove, i.e., the sulcus tali (see below). Like *H. floridanus*, the facets are highly asymmetrical, but the condition in *S. hondaensis* is even more exaggerated, as the ectal facet is much larger than the sustentacular facet. As in the astragalus described above, the ectal facet is kidney-shaped and the sustentacular facet is sub-oval. The ectal facet is located at an oblique angle with respect to the long axis of the tuber calcanei, unlike that of *H. floridanus*. Like other pampatheriids, the calcaneal sustentacular facet of *S. hondaensis* is located anterior to the anterior border of the ectal facet. However, this facet is even more anteriorly placed in *S. hondaensis* than in other pampatheriid species as consequence of its particularly wide sulcus tali. Posteriorly, the calcaneal tuber is not massive in comparison with Pleistocene species of *Holmesina* (e.g., *H. septentrionalis*), but rather mediolaterally compressed, particularly towards its dorsal side.

***Scirrotherium antelucanus*** Laurito and Valerio, 2013

*LSID*. urn:lsid:zoobank.org:act:225CD304-3B63-4B55-B8B8-33B46C90A194

**Holotype.** CFM-2867, mobile osteoderm (Laurito and Valerio, 2013).

**Type locality and horizon.** San Gerardo de Limoncito, County of Coto Brus, Province of Puntarenas, Costa Rica. Upper Curré Formation, Upper Miocene (Laurito and Valerio). For further information about the stratigraphic position of the Curré Formation, see these references: Lowery 1982; Yuan 1984; Rivier 1985; Kolarsky et al. 1995; Alvarado et al. 2009; Aguilar et al. 2010; Obando 2011. There are no published absolute ages for this geological unit.

**Diagnosis.** Unmodified (see Laurito and Valerio, 2013; p. 47).

**Referred material.** MUN STRI 36880, an isolated fixed osteoderm (Fig. 8).

**Figure 8.**
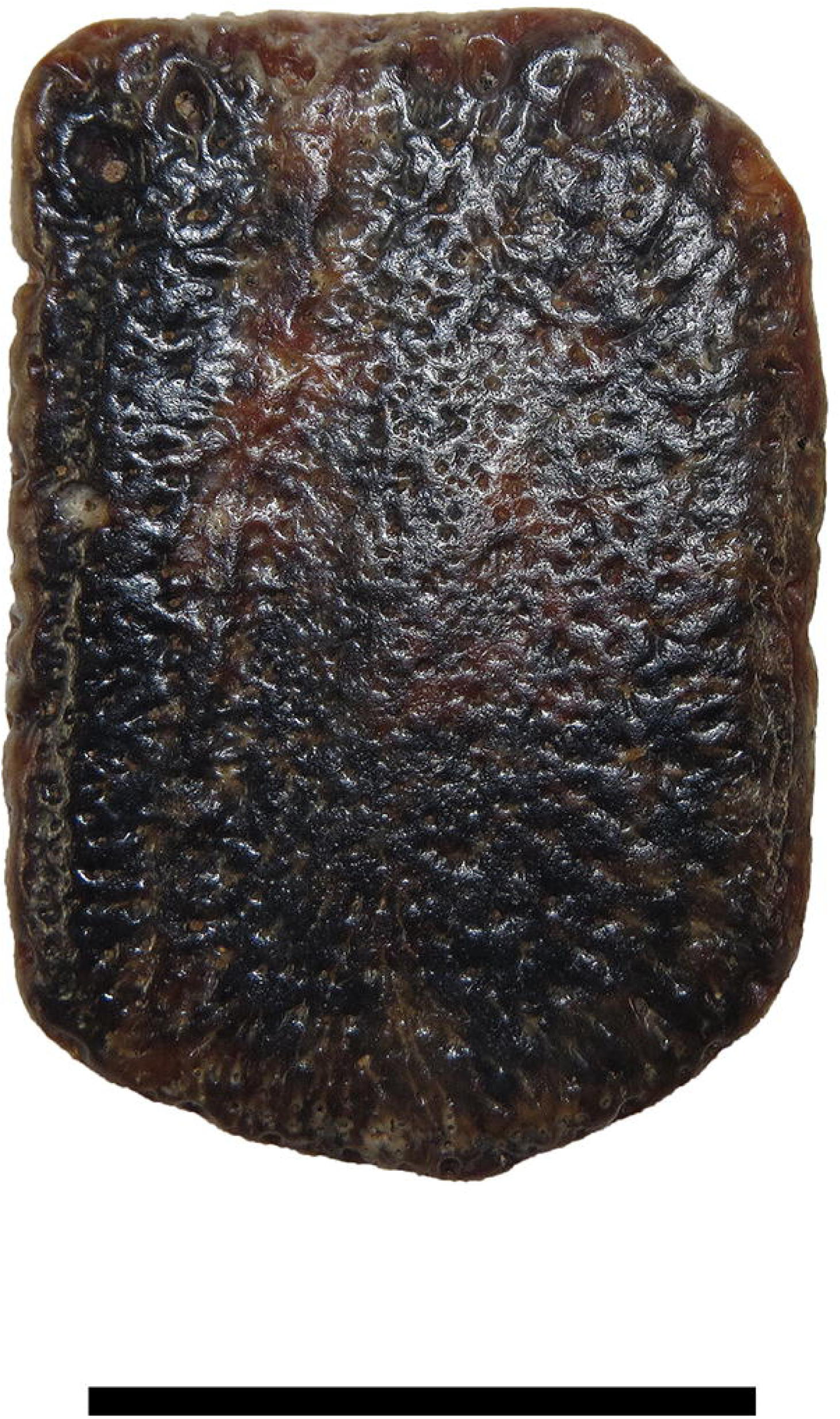
Fixed osteoderm MUN STRI 36880 of *Scirrotherium antelucanus* from the upper Sincelejo Formation, Department of Sucre, Colombia. Scale bar equal to 20 mm.

**Occurrence.** Upper Sincelejo Formation, Upper Miocene to Pliocene (Messinian to Zanclean). El Coley Town, Municipality of Los Palmitos, Department of Sucre, Colombia. For further information about the stratigraphic position of the Sincelejo Formation, see these references: Flinch 2003; Villarroel and Clavijo 2005; Bermúdez et al. 2009; and Alfaro and Holz 2014. There are no published absolute ages for this geological unit.

**Remarks.** The fixed osteoderm MUN STRI 36880, possibly from the of the pelvic shield, is assigned to the species *S. antelucanus* on the basis of the following observations: (I) the area and thickness of this osteoderm (linear measurements in millimetres: anteroposterior length = 34.91; transverse width = 24; thickness = 4.45; approximate area = 837.8 mm^2^) are within the range of variability for comparable osteoderms of *S. antelucanus* and exceed the known values of area for most of the same kind of osteoderms for *S. hondaensis*; (II) the external surface is relatively smooth; (III) the anterior margin is wide; (IV) the anterior foramina are larger (2–3 millimetres of diameter) than in *S. hondaensis*, like *S. antelucanus* from Costa Rica; (V) the number of anterior foramina (9) is within the range of variability for *S. antelucanus* (7–10 for quadrangular osteoderms, like the specimen here described), but greater than the range for *S. hondaensis*; (VI) poorly elevated longitudinal central elevation, like in some osteoderms of *S. antelucanus* (the longitudinal central elevation is generally more elevated in *S. hondaensis*; see Laurito and Valerio 2013).

### aff. Scirrotherium

**Referred material**: MUN STRI 16718 (Fig. 9(A)), fixed osteoderm of the scapular shield; MUN STRI 38064 (Fig. 9(E)), undetermined fixed osteoderm; MUN STRI 16719 (Fig. 9(G)), mobile osteoderm fragmented in its anterior margin.

**Figure 9.**
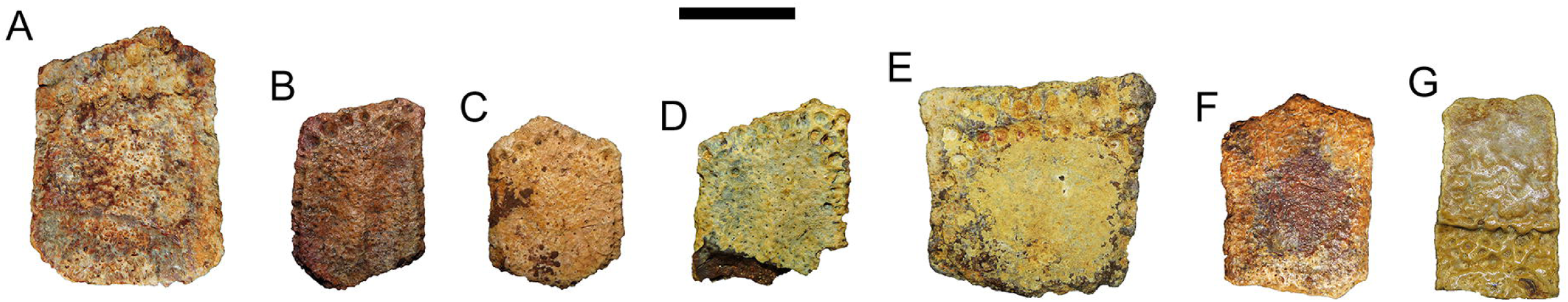
Pampatheriid osteoderms from the Department of La Guajira, Colombia, referred to as aff. *Scirrotherium* (**A**, MUN STRI 16718; **E**, MUN STRI 38064; and **G,** MUN STRI 16719; all these specimens are from the Castilletes Formation and they are fixed osteoderms except the latter, which consist of an anterior fragment of a mobile osteoderm); *Scirrotherium* cf. *hondaensis* (**C**, MUN STRI 36814, a fixed osteoderm from the Castilletes Formation); and *Scirrotherium* sp. (**B**, MUN STRI 36801; **D**, MUN STRI 16158; and **F**, MUN STRI 34373; all these fixed osteoderms are from the Castilletes Formation, except the latter, which comes from the Ware Formation). Note the two well-developed rows of anterior foramina in the osteoderms MUN STRI 16718 and 38064. Scale bar equal to 20 mm.

**Occurrence**: Castilletes Formation, upper Lower Miocene to lower Middle Miocene, upper Burdigalian to Langhian). Localities of Makaraipao, Kaitamana and Patajau Valley (localities with numbers 390093, 430202 and 390094 in Moreno et al. 2015, respectively), Municipality of Uribia, Department of La Guajira, Colombia.

**Description**: The fixed osteoderm of the scapular shield MUN STRI 16718 (Fig. 9(A)) is relatively large and has a pentagonal outline. Its linear measurements in millimetres are: anteroposterior length = 45.02; transverse width = 33.41; thickness = 6.66. These values imply that this osteoderm has greater area than any other known area size for osteoderms referred to as *Scirrotherium* (Table 1), including those of the osteoderms of the larger *Scirrotherium* species, i.e., *S. antelucanus* (see Appendix 1 in Laurito and Valerio 2013). Rather, this osteoderm is similar in size to those reported for *H. floridanus*. The external surface of the osteoderm MUN STRI 16718 is punctuated by numerous diminutive pits, like *S. hondaensis* and *S. antelucanus*. The surface does not have a recognizable longitudinal central elevation nor longitudinal depressions, so that the osteoderm has a flattened appearance, similar to that of several osteoderms of *S. antelucanus* (Laurito and Valerio 2013). In contrast, the marginal elevations are easily identifiable. These ridges are relatively low and narrow. There are foramina with a nearly homogeneous large size in the anterior margin. They are aligned in two well defined rows. The most anterior row has five foramina and the posterior one has six. Collectively, the two rows of foramina rows are equivalent to ca. 25% of the anteroposterior length of the osteoderm. In *S. hondaensis*, the rows of foramina in fixed osteoderm, when present, this percentage comprises less than 20%.

The osteoderm MUN STRI 38064 (Fig. 9(E)) does not appears to be a non-marginal osteoderm. It has a trapezoidal outline and the following measurements in millimetres: anteroposterior length = 39.08; transverse width = 39.55; thickness: 5.98. These values are within the range of variability of *S. antelucanus*. This osteoderm has two long rows of anterior foramina in which the posterior row seems to extend partially over the anterior lateral margins, unlike the anterior foramina row(s) in *S. hondaensis* and *S. antelucanus*. The most anterior row of foramina is formed by eight foramina and the posterior row has 11 foramina. In both of these rows, the foramina are of similar size, although a few are smaller. The rows of foramina diverge on the left lateral margin and within the resultant space between these rows is located a large and isolated foramen. This osteoderm does not have a recognizable longitudinal central elevation nor longitudinal depressions, i.e., it is flattened. Its marginal elevations are narrow and poorly elevated. The foramina of the lateral margins are smaller than most of anterior foramina. As a consequence of preservation factors, the pits on the external surface are not present.

The osteoderm MUN STRI 16720 is a partial mobile osteoderm (Fig. 9(G)) with an elongated rectangular shape. Its linear measurements in millimetres are: anteroposterior length (incomplete by fragmentation) = 45.68; transverse width = 30.69; thickness = 6.96. The external surface is convex and without a longitudinal central elevation or longitudinal depressions. The anterior margin shows a set of foramina not clearly aligned in rows.

**Remarks**: With current evidence, the osteoderms MUN STRI 16718 and MUN STRI 38064 could not be confidently assigned to *Scirrotherium*. This taxonomic decision is supported by several arguments. First, morphologically, the osteoderms referred here to aff. *Scirrotherium* are more similar to those of *S. hondaensis* and *S. antelucanus* than to any other osteoderms of known pampatheriids. The osteoderms of aff. *Scirrotherium* differ with respect to the osteoderms of *S. hondaensis* and *S. antelucanus* in three characteristics: (I) a greater number of anterior foramina and/or greater development of two rows from these foramina; (II) longitudinal central elevation possibly absent, i.e., the presence of a flattened external surface; (III) larger maximum osteoderm area. Of these features, the third one (III) is the least ambiguous, i.e., the maximum area of fixed osteoderms exceeds those of the osteoderms of *S. hondaensis* and *S. antelucanus*. Comparatively, the first and second (I and II) characteristics are more ambiguous considering that similar conditions were also observed in *S. hondaensis* and *S. antelucanus*. These conditions are described as follows.

Some infrequent osteoderms of *S. hondaensis* have two anterior rows of foramina, of which the anterior row is comparatively less developed (i.e., with smaller and fewer foramina) than in aff. *Scirrotherium*. Additionally, in *S. hondaensis* and, particularly in *S. antelucanus*, some osteoderms have a flattened or even a missing longitudinal central elevation. These observations imply limitations on the taxonomic resolution, especially considering that the material on which aff. *Scirrotherium* is based is scarce and does not allow comparisons of a representative series of osteoderms encapsulating the morphological variability within the carapace of this animal.

***Kraglievichia*** Castellanos, 1927

*LSDI.* urn:lsid:zoobank.org:act:92C8B169-4F79-467E-B951-EF1DE6E327B1

**Type species**: *Kraglievichia paranensis* Ameghino, 1883

**Other referred species**: *Kraglievichia carinatum* comb. nov. (= *Scirrotherium carinatum*

Góis, Scillato-Yané, Carlini and Guilherme, 2013; see below)

**Emended differential diagnosis**: Small-to-middle sized pampatheriid characterized by fixed osteoderms with ornamentation (particularly the longitudinal central elevation) more conspicuous than in any other pampatheriid; anteriorly wide and posteriorly tapered longitudinal central elevation; very deep longitudinal depressions; highly elevated and frequently blunt marginal elevations, even flattened towards their top; external surface of osteoderms generally rougher than in *Scirrotherium* but less than in *Holmesina*.

***Kraglievichia carinatum*** comb. nov.

2013. *Scirrotherium carinatum –* Góis, Scillato-Yané, Carlini and Guilherme, Fig. 4.

**Holotype**: MLP 69-IX-8-13-AB, a mobile osteoderm (Góis et al. 2013).

**Type locality and horizon**: Paraná River cliffs, Province of Entre Ríos, Argentina. Ituzaingó Formation, Upper Miocene, Tortonian (Góis et al. 2013).

**Differential diagnosis**: Unmodified (see Góis et al. 2013, p. 182).

**Referred material**: The holotype, paratypes and part of the hypodigm of this species (see Fig. 10 and Appendix S1 of the Supplementary Material).

**Figure 10.**
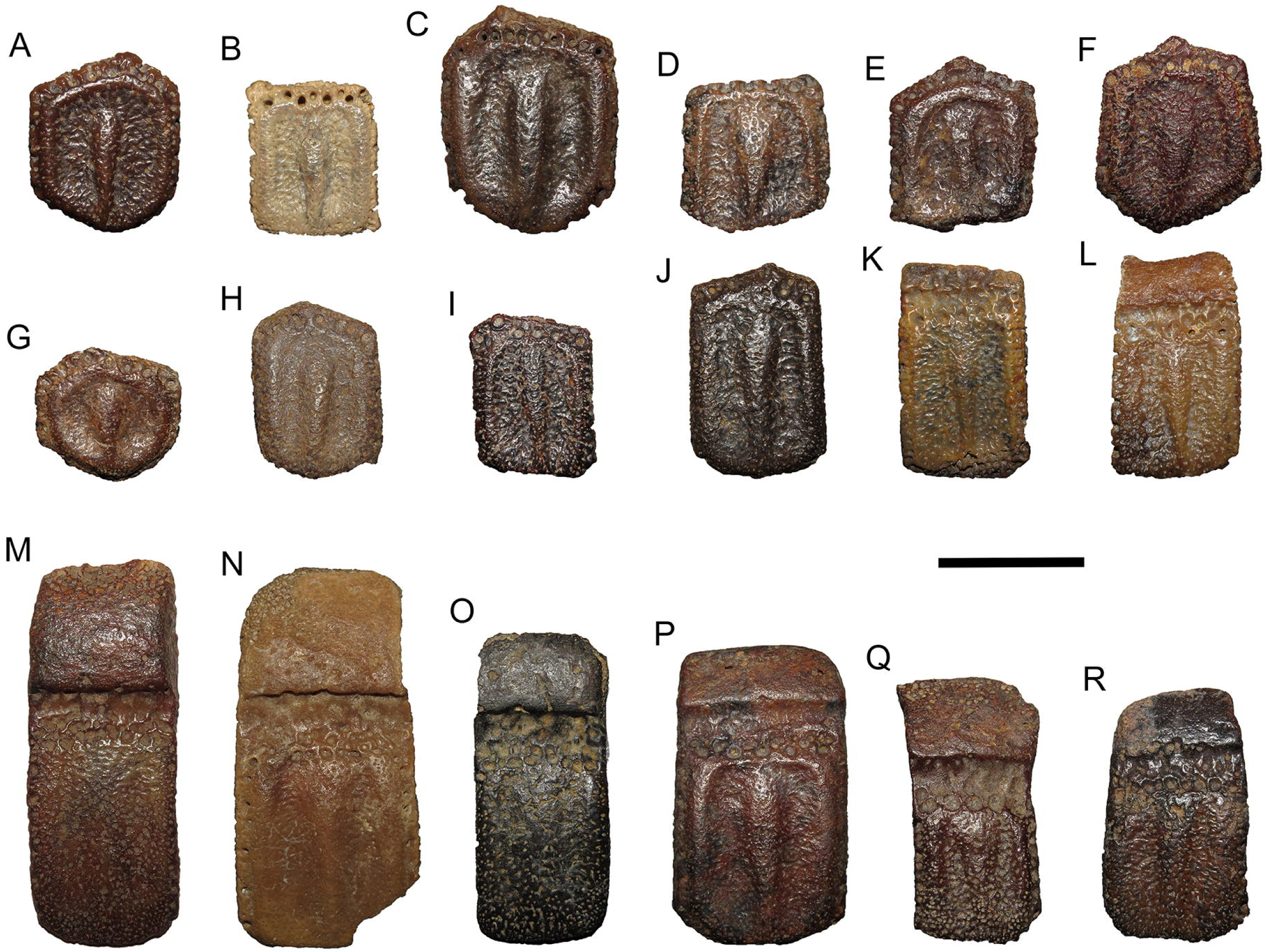
Osteoderms of *Kraglievichia carinatum* comb. nov. from the Ituzaingó Formation, Province of Entre Ríos, Argentina. The holotype of this species is marked with one single asterisk (*) and paratypes with double asterisk (**). A–J, fixed osteoderms; K– R, (semi) mobile osteoderms. **A**, MLP 69-IX-8-13AC**; **B**, MLP 70-XII-29-1**; **C**, MLP 41-XII-13-905; **D**, MLP 69-IX-8-13AF; **E**, MLP 69-IX-8-13AG; **F**, MLP 41-XII-13-414A; **G**, MLP 69-IX-8-13AN; **H**, unknown catalog number; **I**, MLP 69-IX-8-13AK; **J**, MLP 41-XII-13-414B; **K**, MLP 69-IX-8-13AS; **L**, MLP 69-IX-8-13AE**; **M**, MLP 52-X-1-36; **N**, MLP 69-IX-8-13AB*; **O**, MLP 41-XII-13-909; **P**, MLP 69-IX-8-13AW; **Q**, MLP 69-IX-8-13AQ; **R,** MLP 69-IX-8-13AY. Scale bar equal to 20 mm.

**Discussion.** In their descriptive work on *K. carinatum* comb. nov., Góis et al. (2013) did not explicitly justify the inclusion of this species within *Scirrotherium*. Interestingly, part of the material assigned to the taxon they create, coming from northwestern Brazil (Solimões Formation), was previously referred to as *Kraglievichia* sp. by several researchers, including Góis himself (Góis et al. 2004; Góis 2005; Cozzuol 2006; Latrubesse et al. 2010; Góis et al. 2013). However, Góis et al. (2013) refuted the original taxonomic assignment, arguing that it was erroneous, although they did not offer any concrete support for their decision. In the absence of a phylogenetic analysis in Góis et al. (2013), we could assume by default that these authors included *K. carinatum* comb. nov. within *Scirrotherium* because they considered osteoderm features of this species at least compatible with the generic diagnosis proposed by Edmund and Theodor (1997). Based on morphological similarity, Góis and colleagues hypothesized closer affinities between *K. carinatum* comb. nov. and *S. hondaensis* than those between *K. carinatum* comb. nov. and *K. paranensis*. First, it is necessary to analyse in detail each of the osteoderm features of *K. carinatum* comb. nov. in relation to the original diagnosis of *Scirrotherium*. According to Edmund and Theodor (1997), the fixed osteoderms of *Scirrotherium* have a small (not specified) number of large piliferous foramina on the anterior margin. These foramina are well spaced but interconnected by a distinct channel. This is observed both in *K. carinatum* comb. nov. and *S. hondaensis*. Likewise, the presence of continuous marginal elevations, posteriorly confluent with the longitudinal central elevation, is a trait also shared by the two compared species. Finally, the relative osteoderm size of *K. carinatum* comb. nov. is small relative to other pampatheriids, which is in line with the original diagnosis for *Scirrotherium*.

Therefore, the osteoderm features of *K. carinatum* comb. nov. are compatible with those mentioned in the diagnosis for *Scirrotherium* by Edmund and Theodor (1997). However, this does not necessarily imply that the taxonomic allocation of *K. carinatum* comb. nov. to the genus *Scirrotherium* is correct. In fact, there are several reasons to consider this assignment is unreliable. Initially, the diagnosis of Edmund and Theodor (1997) contained only three allegedly diagnostic osteoderm features, including the relative osteoderm size. Furthermore, and more importantly, these “diagnostic features” do not allow definitive discrimination between *Scirrotherium* and any other genus of pampatheriids. Indeed, in their analysis, Góis et al. (2013) accept that, for instance, *Vassallia minuta* also shares the presence of fixed osteoderms with a small number of large anterior foramina, which are well spaced and connected by a canal. Independently, Laurito and Valerio (2013) also highlighted the non-diagnostic nature for *Scirrotherium* of the latter osteoderm trait. The other osteoderm features under consideration, i.e., the posterior confluence of the marginal elevations with the longitudinal central elevation and the small osteoderm size, are also of ambiguous diagnostic value. For instance, the confluence of marginal elevations and longitudinal central elevation is also found in *K. paranensis*, a pampatheriid clearly different from *Scirrotherium*. And although apparently informative on body-size trends of some individual pampatheriid lineages (e.g., *Holmesina* spp.) and useful as discriminant factor between species (Góis et al. 2013; Laurito and Valerio 2013), the relative osteoderm size is not necessarily sufficient to make taxonomic assignments to the genus level among pampatheriids as a whole (see below). In this sense, it is worth noting the potentially conflicting taxonomic conclusions that could be reached using osteoderm-inferred relative body size versus those based on non-osteoderm evidence. For example, MLP 69-IX-8-13A, a femur belonging to an adult individual from the Ituzaingó Formation, was assigned to *K.* cf. *paranensis* by Scillato-Yané et al. (2013), is comparable in size to that of the small pampatheriid *S. hondaensis*. Therefore, it is probable that the referred femur does not belong to the medium-to-large sized *K. paranensis*, although it is reasonable to include it in the genus *Kraglievichia* (as the authors decided). However, Scillato-Yané et al. (2013) did not discuss the possibility that the material assigned to *K.* cf. *paranensis*, particularly MLP 69-IX-8-13A, might be related to the (partially) co-occurrent species of *K. paranensis*, i.e., *K. carinatum* comb. nov., a pampatheriid whose small body size is fully compatible with the small size of that femur. In other words, like Góis et al. (2013), they did not seriously consider the hypothesis of *K. carinatum* comb. nov. as a small species of *Kraglievichia*, rather than a species belonging to *Scirrotherium*.

Again analysing the original diagnosis of *Scirrotherium* by Edmund and Theodor (1997), it should be regarded as ambiguous and hardly useful to differentiate this genus from other genera in Pampatheriidae, at least with respect to osteoderm traits. It is likely that these supposedly diagnostic features are actually symplesiomorphies for the entire family or, at most, a hypothetical subfamilial lineage. This means that Góis et al. (2013) did not have a sufficiently robust diagnosis of *Scirrotherium* to confidently assign *K. carinatum* comb. nov. to this genus. Alternatively, they may have noted morphological similarity between osteoderms of *K. carinatum* comb. nov. and *S. hondaensis* from features not included in the diagnosis by Edmund and Theodor (1997). However, Góis et al. (2013) only listed morphological differences between these species and virtually did not mention any similarity for them, except for potentially equivocal resemblance as that indicated by relative osteoderm size (i.e., small osteoderm sizes in comparison with those of *K. paranensis* and *Plaina*). The lack of usefulness of the relative osteoderm size for generic assignation is further supported by the osteoderm morphometric analysis in Góis et al. (2013, p. 185), whose resulting PCA and CCA plots show that, despite the similarity in relative osteoderm size, *K. carinatum* comb. nov. is located far from *S. hondaensis* (which is closer to *V. minuta*, a taxon apparently related to other main lineage within Pampatheriidae, i.e., *Plaina*-*Pampatherium*) and *K. paranensis* in morphospace.

Summarizing, there is little justification by Góis et al. (2013) on their taxonomic decision to including *K. carinatum* comb. nov. within *Scirrotherium*. Considering the morphological conservatism of Pampatheriidae, the observation of a common general morphological pattern between *K. carinatum* comb. nov. and *S. hondaensis* does not necessarily imply the grouping of these species under the same generic taxon, least of all by omitting the taxonomic significance of striking similarities in osteoderm ornamentation between *K. carinatum* comb. nov. and the better known taxon *K. paranensis*. Furthermore, we should note that Góis and colleagues, in their work on *K. carinatum* comb. nov., did not make morphological comparisons including to *S. antelucanus*, a species more similar in osteoderm features to *S. hondaensis* (i.e., the type species of *Scirrotherium*). The species *S. antelucanus* was described on a scientific article (Laurito and Valerio 2013) published nearly simultaneously, but later, to that of *K. carinatum* comb. nov. Therefore, Góis et al. (2013) did not know about the existence of *S. antelucanus* when they performed their systematic analysis (“Until the present study, *S. hondaensis* was the only known species of this genus”; Góis et al. 2013, p. 177), so that their taxonomic assignment of *K. carinatum* comb. nov. to *Scirrotherium* was likely the result of limited notions on the morphological variability and diversity of *Scirrotherium* in northern South America and southern Central America.

In this work I decide to assign *K. carinatum* comb. nov. to the genus *Kraglievichia* based on the results of phylogenetic analyses that I designed considering the hypothesis of Edmund (1985) on the probable subfamilial relationships within Pampatheriidae, which implicitly sustains that the creation of supraspecific taxa from osteoderm evidence should be determined –with the prerequisite of morphological similarity- by the degree of development of the ornamentation. Understanding that *K. carinatum* comb. nov. has morphologically similar osteoderms to those of *K. paranensis* (apart from relative osteoderm size) and has one of the more conspicuous, protuberant osteoderm ornamentations among pampatheriids, along with *K. paranensis*, as acknowledged by Góis et al. (2013) themselves, this means that *K. carinatum* comb. should be considered closely related to *K. paranensis* and therefore they both should also be included in the same genus, i.e., *Kraglievichia*.

## 5. Discussion

### 5.1. Systematic implications

This systematic analysis is the first attempt to test the intergeneric relationships and internal structure of the genus *Scirrotherium* with its three previously referred species, i.e., *S. hondaensis* (type species), ‘*S.*’ *carinatum* (= *K. carinatum* comb. nov.) and *S. antelucanus*. The two strict consensus trees from the distinct character weighting schemes (equal and implied weights) show very similar results. However, the preferred phylogenetic hypothesis is that supported by the implied weights analysis. According to Goloboff et al. (2018), the parsimony analysis under implied weights outperforms equal weighting and the model-based methods. Beyond this preference for a particular hypothesis (further supported below), both resultant trees agree that the species referred to as *Scirrotherium* are not monophyletic and, consistently, from a diagnostic point of view only *S. hondaensis* and *S. antelucanus* appears as those actually referable to *Scirrotherium*. The relationship between *S. hondaensis* and *S. antelucanus* could not be confidently resolved, despite the inclusion of new osteoderm characters (the only ones comparable between these species so far) in these parsimony analyses. Consequently, a paraphyletic relationship among *S. hondaensis* and *S. antelucanus* should not be rule out. In conjunction, these results suggest the need of information on craniomandibular or dental characters for *S. antelucanus* to further test the affinities of this species with respect to *S. hondaensis*. Until new phylogenetic evidence is available, *Scirrotherium* is maintained as taxonomically valid using a new, emended diagnosis which focus (in addition to specific osteoderm similarities between *S. hondaensis* and *S. antelucanus*) on its lesser degree of development of the osteoderm ornamentation in comparison with those in *Holmesina* and, particularly, *Kraglievichia*, but greater degree of development than in the *Plaina-Pampatherium* lineage. This new diagnosis replaces the original and now inadequate diagnosis by Edmund and Theodor (1997).

Unlike the unpublished phylogeny of Góis (2013), the phylogenetic position of ‘*S.*’ *carinatum* was resolved here, i.e., this species is the sister taxon of *K. paranensis*. Therefore, it is proposed the new name *K. carinatum* comb. nov. Despite Góis (2013) did not recover as a clade to *S. hondaensis* and *K. carinatum* comb. nov., as expected if both these species were assigned to *Scirrotherium*, he presented one supposed synapomorphy that join them, i.e., very deep longitudinal depressions, “in particular in *S. carinatum*” (Góis 2013, p. 215). This feature is not a confident support for grouping *S. hondaensis* and *K. carinatum* comb. nov. because the deepest longitudinal depressions in this pampatheriid lineage are found in *K. carinatum* comb. nov. and *K. paranensis*, not in *S. hondaensis*. The new emended differential diagnosis for *Kraglievichia* acknowledges the highly sculpted external osteoderm surface documented on this taxon, which is more protuberant than in the Plio-Pleistocene genus *Holmesina*. This diagnosis provides an updated and concise description of useful osteoderm features to distinguish *Kraglievichia* from other genera within Pampatheriidae. It is important to note that the species *K. paranensis* has several autapomorphies (see Appendix S4 of the Supplementary Material) which, in addition to the relative osteoderm size, need to be compared in the future with homologous, unknown endoskeletal traits in *K. carinatum* comb. nov. in order to test the phylogenetic affinity of these two species as inferred from osteoderm traits. Provisionally, the difference in relative osteoderm size (consequently also in relative body size) and some morphological differences between *K. carinatum* comb. nov. and *K. paranensis*, as noted by Góis et al. (2013), may be linked to distinct ontogenetic growth trajectories in these species, with *K*. *carinatum* comb. nov. representing the plesiomorphic condition (i.e., small body size; see Sánchez-Villagra 2012 for a discussion on the implications for taxonomy of the ontogenetic growth in extinct species).

Other important difference here with respect to the phylogeny of Góis (2013) is that *K. paranensis* was not recovered as the one single sister taxon of *Holmesina* spp. (except *H. floridanus*). Instead, it is part of a group additionally formed by *S. hondaensis*, *S. antelucanus* and *K. carinatum* comb. nov. Together, these taxa are the sister clade of *Holmesina*. This means that we do not have evidence of direct ancestral forms to *Holmesina* yet. However, despite some striking differences, it is remarkable the greater morphological similarity of the osteoderm ornamentation and several cranial features between *Holmesina* (especially *H. floridanus*) and *Scirrotherium*, rather than with those of *Kraglievichia*. The recognizable similarities between *H. floridanus* and *S. hondaensis*, and at the same time differences with *K. paranensis*, include less protuberant osteoderm ornamentation; occurrence of uniformly narrow longitudinal central elevation in fixed osteoderms; more robust skull; more expanded frontals; the plane of the frontals forming a reflex angle with that of the parietals; less anteroposteriorly elongated upper teeth; and the first two upper molariforms less obliquely oriented with respect to the midline of the hard palate.

Góis (2013) found that *Holmesina* is non-monophyletic due to the phylogenetic position of *H. floridanus* with respect to the other *Holmesina* species. This result coincides with that recovered here from the parsimony analysis with equal weights. *Holmesina floridanus* has several plesiomorphic features in comparison with the remaining *Holmesina* species, e.g., less protuberant ornamentation, less rough external surface of the osteoderms and less dorsally situated basicranium with respect the palatal plane. However, the confluent arrangement of the calcaneal facets of the astragalus in *H. floridanus* suggests that this species might not be directly related to any known South American pampatheriid (Edmund 1987). Likewise, Gaudin and Lyon (2017) have recently found potential support for the monophyly of *Holmesina* from craniomandibular specimens. Therefore, the position of *H. floridanus* in a polytomy with *S. hondaensis* and *S. antelucanus* in the analysis with equal weights is explained from the lack of resolution (relatively small character sampling by a fragmentary fossil record and homoplastic noise), not as product of a “real” non-monophyletic relationship with the remaining *Holmesina* species. Conversely, the *Holmesina* is recovered as monophyletic by the implied weights analysis. As expected, in this topology *H. floridanus* is the most basal species of *Holmesina*. In both strict consensus trees, *H. septentrionalis*, the other North American species, is grouped together with all the South American *Holmesina* species (except *H. rondoniensis* and *H. cryptae*, both excluded in this study).

I abstained from revising the diagnosis of *Holmesina* because this is considered beyond the intended objectives of this work. However, six putative synapomorphies (unambiguous and ambiguous) are proposed (or further supported) for this genus: (1) Anterior and lateral margins with elongated, strong bone projections as radii directed from the external border of the central figure towards the osteoderm borders in non-marginal fixed osteoderms; (2) anteriorly convergent, nearly in contact medial processes of premaxillae; (3) length of nasals greater than 30% of the maximum anteroposterior length of the skull; (4) conspicuous and anteroposteriorly elongated maxillary ridge (Gaudin and Lyon 2017); (5) relatively small lacrimal; and (6) bilobed posterior upper molariforms (Mf5-Mf9) with incipient trilobulation.

### 5.2. Evolutionary and biogeographic implications

*Scirrotherium* is a pampatheriid genus from the Early Miocene-Late Pliocene of northern South America and southern Central America. This taxon, along with *Kraglievichia*, forms the sister group of *Holmesina* (Fig. 11), a pampatheriid probably originated in tropical southern North America (Mexico? see Woodburne 2010). However, based on the osteological comparisons presented above, which are expanded with respect to those of Edmund (1987), the hypothetical South American ancestor or sister taxon of *Holmesina* probably was morphologically generalized, more similar to *Scirrotherium* or aff. *Scirrotherium* than to *Kraglievichia*. This interpretation is in line with that of Edmund (1987), according to which the calcaneo-astragalar articulation of *S. hondaensis* challenges the hypothesis that this pampatheriid is ancestral to *H. floridanus*, but the ornamentation pattern of the osteoderms suggests “at least some degree of relationship” with the latter species. Anatomically, *Kraglievichia* should be considered a highly divergent taxon, especially to taking into account its Miocene age. According to Edmund (1987, p. 16), “the osteoderms [of *H. floridanus*] are quite dissimilar to those of *Kraglievichia*”. This interpretation is shared in this work and contrasts with that of Scillato-Yané et al. (2005), which suggested that *Holmesina* originated from a hypothetical South American basal form of *Holmesina* or *Kraglievichia*. It also is opposed to Simpson (1930) and to the phylogeny of Góis (2013) in which *Kraglievichia* is the only sister taxon of *Holmesina*.

**Figure 11.**
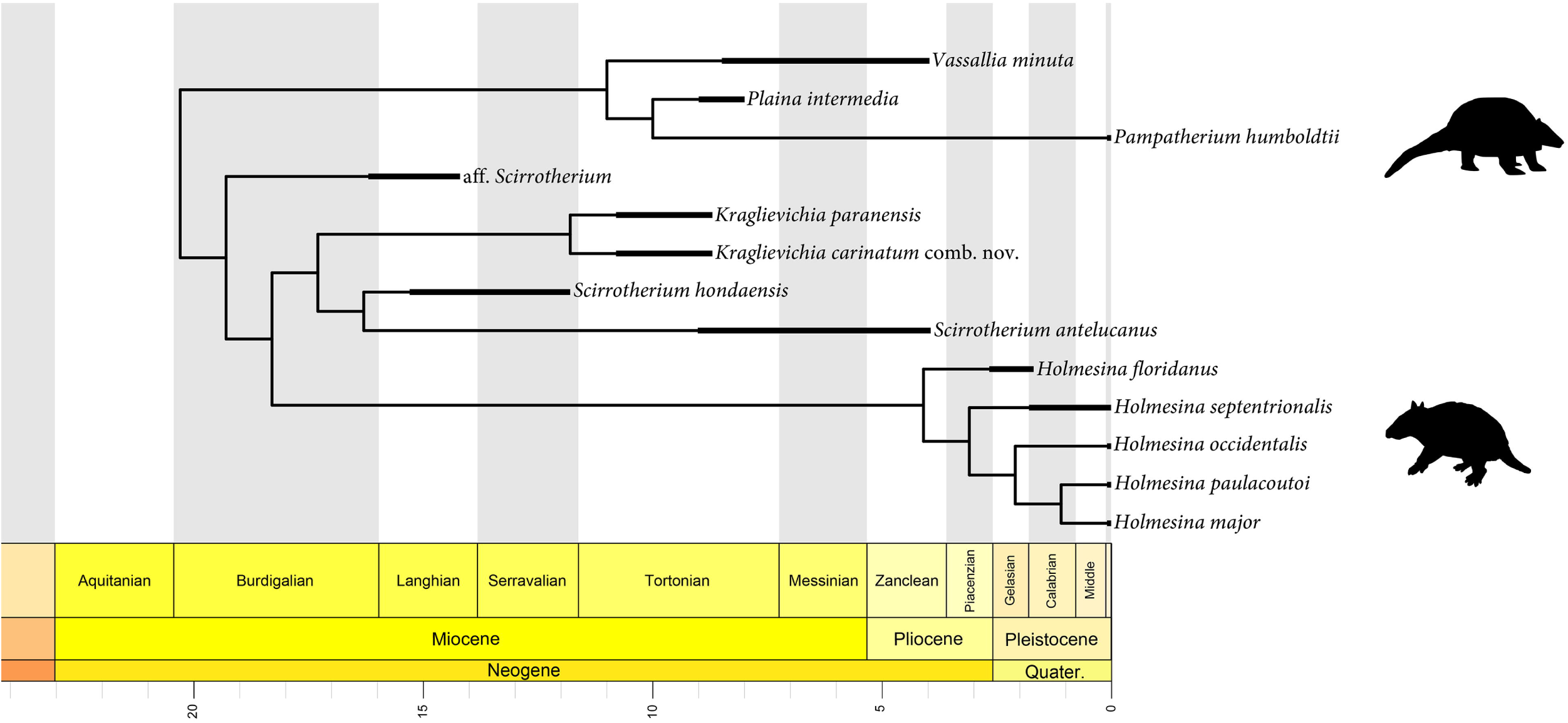
Hypothetical time-calibrated phylogeny of the clade *Scirrotherium* + *Kraglievichia* + *Holmesina* based on the strict consensus tree under implied weights (Fig. 2(B)). Polytomies were resolved by (1) forcing the monophyly of *S. hondaensis* and *S. antelucanus* and (2) placing the species *H. septentrionalis* and *H. occidentalis* as successively basal to the largest South American *Holmesina* species, i.e., *H. paulacoutoi* and *H. major*. Note the diversification events of the clade *Scirrotherium* + *Kraglievichia* + *Holmesina* are mainly concentrated during the Burdigalian (late Early Miocene) and Plio-Pleistocene. Likewise, note the relative long ghost lineage of *Holmesina*. Images of the pampatheriids are from *PhyloPic* (all available under public domain): top, *Pampatherium humboldtii* (http://phylopic.org/name/670230e9-4775-493c-b3ab-31718fb570a3); below, *Holmesina floridanus* (http://phylopic.org/name/73635941-ed8a-4518-aae8-70e824dbee97).

The earliest record of *Scirrotherium*, here treated as tentative by scarce and poorly preserved material, comes from the Early Miocene (late Burdigalian) of northwestern Venezuela (Rincón et al. 2014; see below). Independently from the validity of occurrence of *Scirrotherium* in an Early Miocene locality of northern South America, this record represents the oldest pampatheriid reported in the scientific literature so far. This improvement makes the fossil record of Pampatheriidae more congruent with the expected time of origination of this family, i.e., Late Oligocene-Early Miocene, according to the very few available time-calibrated phylogenies including representatives of Pampatheriidae and its sister group, Glyptodontidae (e.g., Fernícola 2008; Billet et al. 2011). Apparently, the Early Miocene Venezuelan pampatheriid indicates an origin of these xenartrans in low latitudes in South America. However, this hypothesis could be defied by a possible Late Eocene pampatheriid of Argentina, which has been not formally described and published yet (Góis 2013).

Beyond the geographic origin of Pampatheriidae, northern South America seems to have been a critical area for the early diversification of, at least, the lineage including *Scirrotherium*. This genus probably differentiated at least as early as the late Early Miocene-early Middle Miocene (late Burdigalian-Langhian) in northernmost South America. The former evolutionary inference is consistent with the late Early Miocene record referred to as *Scirrotherium* from Venezuela (Rincón et al. 2014).

Collectively, *Scirrotherium* and *Kraglievichia* occupied a large area in South America during the Neogene (Fig. 12). The geographic range of *Scirrotherium* was more restricted than that of *Kraglievichia*, comprising only tropical low latitudes, instead of a wide latitudinal range, as suggested by Góis et al. (2013). The revaluated distributional pattern of *Scirrotherium* is comparable to that of the glyptodontid *Boreostemma*, which is recorded from the Middle Miocene to the Late Pliocene of Colombia and Venezuela (Carlini et al. 2008; Zurita et al. 2016). In contrast, the distributional range of *Kraglievichia* is similar to that of other Miocene xenarthran taxa at the generic and specific level, which occurred in southern South America and northwestern Brazil, but not in the northern or northwestern end of South America (see Ribeiro et al. 2014).

**Figure 12.**
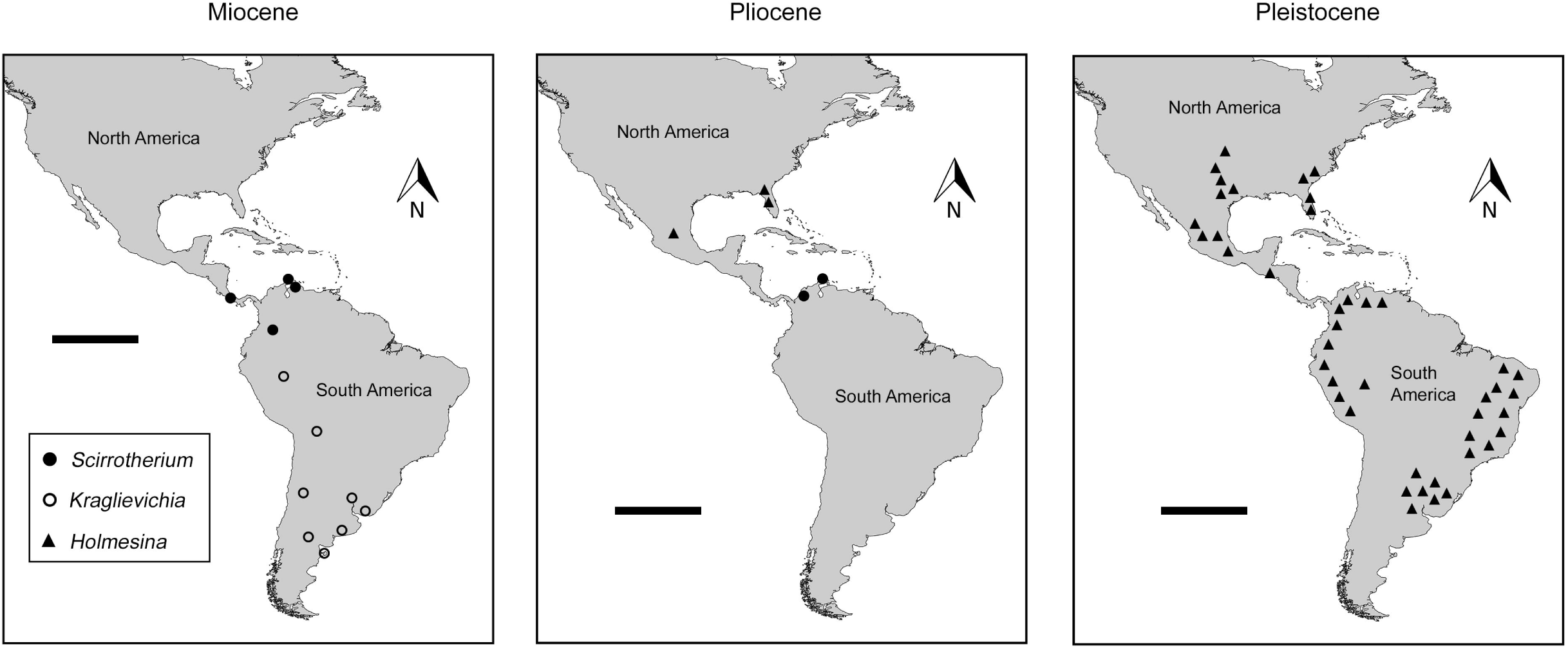
Geographic distributions of *Scirrotherium*, *Kraglievichia* and *Holmesina* during the Neogene and Pleistocene. The symbols (i.e., circles and triangles) should not necessarily be interpreted as single localities but as approximate areas of occurrence. This is especially true for the Pliocene and Pleistocene epochs. Further details on the biogeography of these genera in the *Discussion* section. Scale bar equal to 2000 Km.

Overall, this evidence indicating biogeographic divergence of northwesternmost South America as an independent faunal province from the late Early Miocene to Middle Miocene, and possibly into the Late Miocene or even the Early Pliocene, is consistent with the results of multiple analyses of the South American terrestrial mammal fossil record for the Neogene (Patterson and Pascual 1968; Cozzuol 2006; Ortiz-Jaureguizar and Cladera 2006; Croft 2007; Carrillo et al. 2015; Rincón et al. 2016; Kerber et al. 2017; Brandoni et al. 2019). Apparently, the existence of one or several strong geographic and/or ecoclimatic barriers (e.g., the Pebas Mega-Wetland System, whose expansion climax coincides with the Middle Miocene) in northern South America would explain that regional endemism pattern (MacFadden 2006; Croft 2007; Salas-Gismondi et al. 2015; Jaramillo et al. 2017). At the same time, the development of a late Early-to-Middle Miocene biogeographic divergence between northwesternmost South America and the rest of this continent may account for the evolutionary divergence of *Scirrotherium* and *Kraglievichia*.

In the Late Miocene, without a completely formed Panamanian Land Bridge (O’dea et al. 2016), *Scirrotherium* expanded its geographic range to southern Central America (Fig. 12), suggesting a possible ephemeral land connection or, more likely, overwater dispersal between South America and Central America (maybe via rafting mechanism; efficient active swimming of pampatheriids in a marine channel seems highly improbable). This is the earliest dispersal event of a pampatheriid to North America (see below). The Central American species of *Scirrotherium*, *S. antelucanus*, is larger than *S. hondaensis*, but comparable or even smaller than aff. *Scirrotherium*. From available evidence, it is not possible to determinate the most probable area of evolutionary differentiation of *S. antelucanus*, but now there is support for occurrence of this species in the late Neogene of northwestern South America, specifically in the Department of Sucre, Colombia.

The South American record of *S. antelucanus* is probably several million years younger (3– 5 my) than the Central American record. However, given the lack of absolute dating for the fossil-bearing stratigraphic levels and the occurrence of Late Miocene strata in the same geological unit (Sincelejo Formation) where comes the material here assigned to *S. antelucanus* in Colombia, it should be recognized a significant age uncertainty for the new South American record of this species. In any case, this age is considered may be Early Pliocene or, alternatively, Latest Miocene from the stratigraphic position of the fossil-bearing horizons (Villarroel and Clavijo, 2005; Bermúdez et al. 2009; Alfaro and Holz 2014; Bernal-Olaya et al. 2015; Córtes et al. 2018), as well as from associated palynomorphs (Silva et al. 2012; B. Fernandes and C. Jaramillo, pers. comm. 2014).

The biogeographic correlation across the Isthmus of Panama using *S. antelucanus* has insightful implications for the understanding of the late Cenozoic intercontinental migratory dynamics in the Americas, including the Great American Biotic Interchange (GABI) (Webb 2006; Woodburne et al. 2006; Woodburne 2010; Cione et al. 2015; Bloch et al. 2016).

Noteworthy, this is the first transisthmian biogeographic correlation for a Neogene terrestrial mammal at the level of species; furthermore, it is the first short-distance intercontinental correlation (i.e., adjacent to the Central American Seaway) with high taxonomic resolution for Neogene land mammals of the Americas; and, finally, it constitutes the first evidence of a distributional pattern congruent with a re-entry event to South America by a pre-Pleistocene xenarthran.

We know a few biogeographic correlations across the Isthmus of Panama which are based on records at generic level of Neogene and Pleistocene land mammals, as well as a very few records at species level of the latter epoch. The Neogene biogeographic correlations include the pampatheriid genera *Plaina*, in Mexico and central-southwestern South America; *Pampatherium*, in Mexico and southeastern South America; and *Holmesina*, in the United States, Mexico and El Salvador, as well as in northwestern and southeastern South America (Woodburne 2010). At the level of species, for instance, the Pleistocene megatheriine *Eremotherium laurillardi* has occurrence in both sides of the Isthmus of Panama in North- and South America (Cartelle and De Iuliis 1995, 2006; Tito 2008; McDonald and Lundelius, E. L. Jr. 2009; Martinelli et al. 2012; Cartelle et al. 2015).

The record in South America of *S. antelucanus* increase the taxonomic resolution of transisthmian biogeographic correlations of Neogene land mammals, opening the possibility of new correlations of this kind and their biostratigraphic application in circum-Caribbean basins, in a similar way as envisioned by the renowned American palaeontologist Ruben A. Stirton from his revision of the fossil mammal remains of “La Peñata fauna” (Stirton 1953), the vertebrate fossil association where comes the new record of *S. antelucanus*. This translates into direct correlation of Land Mammal Ages (in this case, SALMA and NALMA) from migrant mammals which are shared at species level by both South and North America. Using to *S. antelucanus*, this would mean exists a support for faunal, not necessarily chronological, correlation of the early Hemphillian and Montehermosan mammal (xenarthran) assemblages in North America and South America, respectively (see Laurito and Valerio, 2013). Naturally, any compelling intercontinental faunal correlation requires more than one taxonomic element for support. The direct intercontinental faunal correlations from Cenozoic land mammals between South and North America are still underdeveloped in comparison with those between other continents (e.g., North America and Europe or North America and Asia; Woodburne and Swisher, C. C. III 1995; Beard and Dawson 1999; Bowen et al. 2002)

Additionally, the transisthmian correlation of *S. antelucanus* allows to increase the geographic resolution in the detection of intercontinental migrations of late Cenozoic land mammals, which are restricted mainly to large and middle distance correlations for the Neogene record (e.g., Mexico-southern South America; Woodburne 2010). This pattern has prevented the exploration of possible early or intermediate phases of anagenetic/cladogenetic events in late Cenozoic Interamerican migrant taxa, which in turn it is reflected in the fact that we are detecting “suddenly” well-differentiated terminal taxa (e.g., *Holmesina*) in marginal, distant areas with respect to the Central American Seaway and adjacent terrains (Cione et al. 2015 and references therein).

On the other hand, the new transisthmian correlation here presented suggests a possible Neogene re-entry event by a xenartran to South America after its evolutionary differentiation in North America (Fig. 12). The confirmation of this depends on a confident determination of the differentiation area for *S. antelucanus*, i.e., if this species originated in South America, the new record is explained more parsimoniously by population maintenance in the ancestral area. Conversely, if this species originated in Central America from a South American species of *Scirrotherium* as *S. hondaensis*, then we are considering a re-entry event to South America. However, as mentioned above, it is not possible to constrain much more than that at this moment. In any case, the possibility of a Neogene re-entry event to South America by a xenarthran is compatible with the fact that we know several of these events during the Pleistocene. Among these Pleistocene events, there are several involved xenarthrans, including the pampatheriids *Holmesina* and *Pampatherium*, the glyptodontid *Glyptotherium*, the pachyarmatheriid *Pachyarmatherium*, the dasypodid *Dasypus* and the megatheriine *Eremotherium* (Woodburne et al. 2006; Woodburne 2010 and references therein).

On another note, the results of this work have evolutionary implications for the genus *Holmesina* and the multiple Interamerican dispersal events of pampatheriids, including that of *Scirrotherium* (discounting the non-confirmed re-entry event to South America). The genus *Holmesina* has its oldest record (*Holmesina* sp.) in sedimentary rocks deposited around the Pliocene-Pleistocene boundary (ca. 2.4 mya) in Florida, United States (Edmund 1987; Woodburne 2010 and references therein; Gaudin and Lyon 2017). This northward dispersal event is part of the earliest phase of the GABI (GABI 1), in which additionally participated other xenartrans as *Dasypus*, *Pachyarmatherium* and *Eremotherium* (Woodburne 2010). Typically, *H. floridanus* has been considered the most basal among the *Holmesina* species (Edmund 1987), as it is supported here. The hypothetical time-calibrated phylogeny introduced in this work for Pampatheriidae (Fig. 11) suggests that exist a long ghost lineage leading to *Holmesina*, from the Early Miocene (Burdigalian) until the Late Pliocene. The improvement of the fossil record in northern South America, Central America and Mexico will allow to advance in the recognition of probable direct ancestral forms for *Holmesina*.

From the above analysis, a probable model of biogeographic evolution of *Holmesina* is as follows (Fig. 12). A hypothetical pampatheriid close to *Holmesina* or even a hypothetical *Holmesina* species basal with respect to *H. floridanus* dispersed to Central America, Mexico and United States during the Pliocene (Early Pliocene according the time-calibrated phylogeny). Once it was established the genus *Holmesina* in North America with *H. floridanus*, the larger species *H. septentrionalis* diverged and differentiated in the Early Pleistocene of southern United States. Later, *H. septentrionalis* expanded southward to Mexico and Central America during the Early-Middle Pleistocene (Aguilar and Laurito 2009). In the Middle or early Late Pleistocene, possibly *H. septentrionalis* colonized South America, where took place an important diversification, which was likely influenced by the Late Pleistocene climatic changes (Scillato-Yané et al. 2005). This diversification gave origin to the species *H. occidentalis*, *H. rondoniensis*, *H. cryptae*, *H. major* and the most robust pampatheriid, *H. paulacoutoi* (Scillato-Yané et al. 2005; Moura et al. 2019).

As inferred from the phylogeny and derived interpretations here presented, the dispersal events of *S. antelucanus* and *H. floridanus* to North America are independent of each other. This means that the number of northward intercontinental dispersal events of pampatheriids during the late Cenozoic actually is at least three, which in chronological order are: (1) genus *Scirrotherium* (Late Miocene); (2) lineage *Plaina*-*Pampatherium* (Early Pliocene); (3) genus *Holmesina* (undetermined Pliocene). From these events, only the latter, based on the fossil record of *H. floridanus*, is included in the GABI. The remaining two events are classified as part of the macroevolutionary invasion “wastebasket” called “Pre-GABI” (literally, ‘before the GABI’; Woodburne et al. 2006; Woodburne 2010; also named by Cione et al. 2015 as “ProtoGABI”). In the lineage *Plaina*-*Pampatherium*, it was differentiated one genus, *Pampatherium*, and at least three species (*P. mexicanum*, *P. typum* and *P. humboldtii*, being the two latter recorded in South America). Meanwhile, the northward dispersal event of *Scirrotherium* seems to give no origin to any other species different to *S. antelucanus*. Only a confirmed southward intercontinental dispersal event of the *Scirrotherium*-*Kraglievichia*-*Holmesina* clade has been well-established, i.e., that of *Holmesina* to South America in the Middle or early Late Pleistocene (Aguilar and Laurito 2009). This event probably is not part of any of the GABI phases of Woodburne (2010) but it appears to be chronologically located between the GABI 2 and 3.

As it has been shown, the study of more abundant and complete pampatheriid material preserved in Neogene geological units of northern South America, in particular, and the current Intertropical region of the Americas, in general, has the potential of provide us more complex and interesting scenarios on the evolution of this glyptodontoid family and, specifically, the genera *Scirrotherium* and *Holmesina*.

## 6. Conclusion

The monophyly of *Scirrotherium* has been tested through parsimony phylogenetic analyses. This taxon is recovered as paraphyletic. ‘*Scirrotherium*’ *carinatum* forms a clade with *Kraglievichia paranensis* and, therefore, here it is proposed the new name *K. carinatum* comb. nov. The remaining referred species to *Scirrotherium*, *S. hondaensis* and *S. antelucanus*, are designed in aphyly. The taxonomic validity of *Scirrotherium*, as defined here, is maintained from diagnostic evidence. *Scirrotherium* is probably the sister taxon of *Kraglievichia*, and these two genera form the sister clade of *Holmesina*. *Scirrotherium* has occurrence from the late Early Miocene to Late Pliocene of northwestern South America (Colombia and Venezuela) and the Late Miocene of southern Central America (Costa Rica). A geographic origin of Pampatheriidae in northernmost South America is suggested from the fossil record of *Scirrotherium* and a new time-calibrated phylogeny. *Scirrotherium* also represents the earliest member of Pampatheriidae which participated in a dispersal event to North America, specifically to the ancient Central American peninsula. This dispersal event happened when the Panamanian Land Bridge was not fully formed yet. The species *S. antelucanus* lived in Central America and northern Colombia during the late Neogene. This is the first Interamerican biogeographic correlation of a Neogene land mammal with high taxonomic resolution, i.e., at the species level. The record of *S. antelucanus* in both sides of the ancient Central American Seaway is compatible with a possible re-entry event of this pampatheriid to South America. In addition, *Scirrotherium* is not probably the South American ancestor of the originally-endemic North American genus *Holmesina*. In contrast with a previous hypothesis which argues that *Holmesina* may have evolved from *Kraglievichia*, here it is suggested that there is no evidence of direct ancestral forms of *Holmesina*, although the unknown South American ancestor of *Holmesina* may be morphologically more similar to *Scirrotherium*.

## Acknowledgements

I am greatly indebted to all curators and managers of the collections visited for revision of specimens during this work, especially to: Andrés Vanegas (Museo de Historia Natural La Tatacoa, Colombia), Aldo Rincón (Museo Mapuka, Universidad del Norte, Colombia), Jorge Moreno-Bernal (Museo Mapuka, Universidad del Norte), Carlos De Gracia (Museo Mapuka, Universidad del Norte), Richard C. Hulbert Jr. (Florida Museum of Natural History, USA), Kenneth Angielczyk (Field Museum, USA), Bill Simpson (Field Museum of Natural History), Patricia Holroyd (University of California Museum of Paleontology, USA) and Marcelo Reguero (Museo de La Plata, Argentina). I am also grateful to Carlos Jaramillo (Smithsonian Tropical Research Institute, Panama) and his field work team from allowing access to fossils they collected in the Department of La Guajira (Colombia). Likewise, special thanks to Antonio Tovar (University of Sucre, Colombia) by inspiration to explore the Sincelejo Formation; to my field assistants during this exploration in the Department of Sucre (Colombia), Oscar Melendrez, Angel Cruz and Juan Pacheco. I would like to thank César Laurito (Instituto Nacional de Aprendizaje de Costa Rica) and Edwin Cadena (Universidad del Rosario, Colombia) for providing photos of osteoderms of the species *S. antelucanus* and one photo of outcrops of the Castilletes Formation, respectively. This work would not have been possible without logistic support and encourage by Alba Lara, Carolay Jiménez and Mónica Montalvo. An anonymous reviewer provided comments that made it possible to improve the manuscript. Financial assistance for this project was provided by several institutions/agencies: CONICET (Internal Doctoral Fellowship); Field Museum of Natural History (Science Visiting Scholarship); Florida Museum of Natural History (International Travel Grant); University of California Museum of Paleontology (Welles Fund); and The Paleontological Society (PalSIRP-Sepkoski Grant).

## Data archiving statement

Data for this study are available in the Dryad Digital Repository: [Intentionally blank] The nomenclatural acts contained in this work are registered in ZooBank:

*LSID*. urn:lsid:zoobank.org:act:313358B5-3B1F-4902-8C2E-BB07CFCBEE18

*LSID.* urn:lsid:zoobank.org:act:E3B83181-91D6-44C8-90C0-BBAACEC2CDEE

*LSID.* urn:lsid:zoobank.org:act:225CD304-3B63-4B55-B8B8-33B46C90A194

*LSID.* urn:lsid:zoobank.org:act:92C8B169-4F79-467E-B951-EF1DE6E327B1

