## Supplementary material for "Systematic revision and redefinition of the genus *Scirrotherium* Edmund and Theodor, 1997 (Cingulata, Pampatheriidae): Implications for the origin of pampatheriids and the evolution of the South American lineage including *Holmesina*"

**Appendices**

**S1.** List of examined specimens or samples (specimens associated under the same catalogue number) of taxa on which this study was focused

**S2.** List of characters for the cladistic analyses

**S3.** Character matrix for the cladistic analyses

**S4.** Synapomorphies supporting nodes and autapomorphies within the strict consensus tree (implied weights)

**S1. List of specimens examined for this study**

Abbreviations in the section “Material and Methods”. See the section “Systematic Palaeontology” for more details on these fossil material. The holotypic material is marked with asterisk (*).

| **Taxon** | **Specimen catalogue number** |
| --- | --- |
| *Scirrotherium hondaensis* | VPPLT 004, 264, 701, 706, 1683 - MT 18; UCMP 37924, 38066, 39846, 40056, 40058, 40201* |
| *Scirrotherium carinatum* | MLP 69-IX-8-13AB*, 69-IX-8AC, 69-IX-8AD, 52-X-1-35, 69-IX-8-13AE, 70-XII-29-1, 41-XII-12-920, 41-XII-13-407, 41-XII-13-414, XII-13-924, 41-XII-13-905, 41-XII-13-909, 41-XII-13-916, 41-XII-13-920, 41-XII-13-923, 41-XII-13-924, 52-X-1-36, 69-IX-8-13AF-Z |
| *Scirrotherium antelucanus* | CMF-2867*, 1849, 3543, 3546, 2559; MUN STRI 36880 |
| *Kraglievichia paranensis*  or *K.* cf. *paranensis* | MACN Pv 2617, 8968; MLP 69-IX-8-13A, 69-IX-8-13. |
| *Holmesina floridanus* | UF 191448, 125356, 125397, 125354, 125384, 125383, 125367, 125365, 125357, 125376, 125359, 125375, 125363, 125396, 125380, 125374, 125369, 125370, 125382, 125393, 125364, 125366, 125361, 125368, 125381, 125358, 125355, 125377, 125379, 125378, 125388, 125362, 125387, 125339, 125315, 125320, 125351, 125326, 125328, 125329, 125321, 125323, 125381, 125317, 125348, 125318, 125335, 125319, 125360, 125330, 125316, 125324, 125352, 125239, 125237, 125267, 125259, 125230, 125248, 125255, 125264, 125261, 125247, 125236, 125235, 24915, 24918, 24919, 125228, 24932 |
| *Holmesina occidentalis* | ROM 3881, MCL 6063 |
| *Holmesina major* | UZM 2314, MPG-PV 3318 |
| *Holmesina paulacoutoi* | MCL-501/01*, MCL-501/110-126* |
| *Vassallia minuta* | MLP 29-IV-15-4*, 69-XII-26-17 |
| *Plaina intermedia* (labeled as *Vassallia maxima*) | FMNH P14424 |
| *Pampatherium humboldtii* | MHD-P-28, MLP-81-X-30-1 |

**S2. List of characters for the cladistic analyses**

All the following characters are referred to cranial, dental or osteodermal features. All the characters are unordered.

Character 1: External surface of non-marginal fixed osteoderms: (0) smooth external surface by sparse tiny pits, without macroscopically visible fiber bundles; (1) moderately-rough external surface by concentrated pits and/or smoothly, macroscopically visible fiber bundles; (2) very rough external surface by highly concentrated pits and coarse, macroscopically visible fiber bundles.

Character 2. Arrangement and size of the anterior foramina of fixed osteoderms: (0) one single row of small foramina; (1) one single, transversely elongated row of large foramina; (2) two well-defined and transversely elongated rows of large foramina.

Character 3: Depth of longitudinal depressions of the fixed osteoderms: (0) apparently absent; (1) (very) shallow, with a gentle slope towards the marginal elevations; (2) deep, forming an angle close to 90° with the plane of the marginal elevations.

Character 4: Breadth of the longitudinal central elevation of fixed osteoderms: (0) (nearly) absent or broadly diffused longitudinal central elevation; (1) sharply, narrow longitudinal central elevation with anteroposteriorly uniform width; (2) anteriorly wide elevation, strongly tapered towards the posterior end; (3) variable anteroposterior width, frequently with long-pyriform or long-tuber shape which is tapered towards one of its ends.

Character 5: Marginal elevations of fixed osteoderms: (0) scarcely elevated, wide and flattened; (1) well elevated and sharp; (2) highly elevated and frequently blunt, even flattened towards their top.

Character 6: Anterior and lateral margins with elongated, strong bone projections as radii directed from the external border of the central figure towards the borders of non-marginal fixed osteoderms: (0) absent; (1) present.

Character 7: Lateral outline of the rostrum in dorsal view: (0) gradually concave; (1) straight to slightly convex.

Character 8: Degree of convergence of the medial processes of premaxillae: (0) anteriorly convergent, but separated by a relatively wide space between premaxillary bones; (1) anteriorly convergent, nearly in contact.

Character 9: Relative length of nasals (Gaudin & Lyon 2017): (0) less than or equal to 30% of the maximum anteroposterior length of the skull; (1) greater than 30% anteroposterior length of the skull.

Character 10: Conspicuous and anteroposteriorly elongated lateral maxillary ridge (Gaudin & Lyon 2017): absent; present.

Character 11: Depth and position of the infraorbital foramen: (0) shallow and (nearly) at the level of the anterior root of the zygomatic arch; (1) deep and clearly separated and anterior to the anterior root of the zygomatic arch.

Character 12: Development of the lacrimal: (0) large, with the facial process more similar to a parallelogram than to a triangle; (1) small, with the facial process more similar to a triangle than to a parallelogram.

Character 13: Anterior root of the zygomatic arch in ventral view (Gaudin & Lyon 2017): (0) posterolaterally projected respect to the main body of the maxilla; (1) laterally projected respect to the main body of the maxilla in a right angle.

Character 14: Shape of the frontals in lateral view: (0) protruding convexity, posterior to the position of insertion of the anterior root of the zygomatic arch; the plane of the nasals forming a reflex angle with that of the parietals; (1) protruding convexity, at the same level of the insertion of the anterior root of the zygomatic arch; the plane of the nasals forming a reflex angle with that of the parietals; (2) gentle convexity, posterior to the insertion of the anterior root of the zygomatic arch; the plane of the nasals forming a (nearly) straight angle with that of the parietals.

Character 15: Lateral extent of the maxilla with respect to the upper teeth rows: (0) maxilla not laterally expanded, forming a labial margin respect to the teeth row less wide than the width of the fourth upper molariform (Mf4); (1) scarce maxillary lateral expansion, forming a labial margin respect to the teeth row equal or slightly wider than the width of the fourth upper molariform (Mf4); (2) very-laterally expanded maxilla, forming a labial margin respect to the teeth row at least twice as width as the fourth upper molariform (Mf4).

Character 16: Interdental spaces of upper molariforms: (0) very narrow, the alveoli of adjacent teeth are nearly in contact between them; (1) narrow, the alveoli of adjacent teeth are relatively well separated.

Character 17: Partial overlapping of the mesiodistal position of the first three molariforms: (0) absent; (1) present.

Character 18: Lobulation of the fourth upper molariform (Simpson 1930, Castellanos 1937): (0) absent; (1) present (at least incipient).

Character 19: Sixth upper molariform (Mf6) with a posterior lobe clearly lateralized: (0) absent; (1) present.

Character 20: Transverse width of sixth molariform (Mf6) respect to its anteroposterior length: (0) greater or equal to 51%; (1) less than 51% and greater or equal to 41%; (2) less than 41% and greater or equal to 31%; (3) less than 31%.

Character 21: Lobulation of the posterior upper molariforms (Mf5-Mf9) (Simpson 1930; Castellanos 1937; Gaudin & Lyon 2017): (0) bilobed; (1) bilobed with incipient trilobulation; (2) trilobed.

Character 22: Lateral expansion of palatines with respect to the ventroposterior region of the maxilla: (0) absent; (1) present.

Character 23: Relative postpalatal length: (0) less than 24% of the greatest palatal length; (1) equal to or greater than 24% and less than or equal to 30% of the greatest palatal length; (2) greater than 30% of the greatest palatal length.

Character 24: Cranial constriction in the orbital-postorbital region in dorsal view: (0) uniformly curved and deeply concave orbital-postorbital outline; (1) orbital-postorbital outline forming a cavity from a nearly straight or poorly curved outline.

Character 25: Relative dorsoventral position of the occipital condyles: (0) at the same level or ventral to the palatal plane; (1) dorsal to the palatal plane.

Character 26: Posterior extent of the parietals and the nuchal crest in lateral view: (0) the posterior border of the nuchal crest is located anterior or at the same level of the posterior border of the occipital condyles; (1) the posterior border of the nuchal crest is located posterior to the posterior border of the occipital condyles.

Character 27: Lobulation of the last lower molariform: (0) incipiently bilobed; (1) bilobed.

**S3. Character matrix used for the cladistic analyses**

This matrix was analyzed using PAUP version 4.0a142. The character state for the character 23 in *S. hondaensis* was estimated.

*Scirrotherium antelucanus*	111110?????????????????????

*Scirrotherium hondaensis*	1111100000100000000000?0??0

*Scirrotherium carinatum*	112220?????????????????????

*Holmesina septentrionalis*	211311??1111????01??1???111

*Holmesina major*	2113110?111?110101??1??011?

*Holmesina paulacoutoi*	2113110?11111101?10?11?011?

*Holmesina occidentalis*	21131101111111010102110011?

*Holmesina floridanus*	111111011111110001011020011

*Kraglievichia paranensis*	112220100010120011000?20000

*Pampatherium humboldtii*	000000201001112101132001111

*Plaina intermedia*	000000011001111001122011111

*Vassallia minuta*	000000????????????????????1

Castilletes specimens	120010?????????????????????

**S4. Synapomorphies and autapomorphies within the strict consensus tree (implied weights)**

Numbers out of parentheses are character numbers and numbers in parentheses are state characters. Only apomorphies of ingroup taxa are listed. Ambiguous synapomorphies are shown in bold.

| **Node/species** | **Synapomorphies/autapomorphies** |
| --- | --- |
| Castilletes specimens | 2 (2) |
| *S. hondaensis* + *S. carinatum* + *S. antelucanus K. paranensis* +  *Holmesina* spp. | 3 (1), 4 (1), 11 (1) |
| *S. hondaensis* | 13 (0), 14 (0), 18 (0), **27 (0)** |
| *S. carinatum + K. paranensis* | 3 (2), 4 (2), 5 (2) |
| *K. paranensis* | 7 (1), 14 (2), 17 (1), **23 (2)**, 26 (0), **27 (0)** |
| *Holmesina* spp. | 6 (1), **8 (1)**, **9 (1)**, 10 (1), **12 (1)**, 21 (1) |
| *H. floridanus* | 20 (1), **23 (2)** |
| *H. septentrionalis* + *H. occidentalis* + *H. major* + *H. paulacoutoi* | 1 (2), 4 (3), **16 (1)**, 22 (1), **25 (1)** |
| *H. occidentalis* | **20 (2)** |
